# Global distributions and emergence of six major tropical root-knot nematodes

**DOI:** 10.64898/2026.02.25.708007

**Authors:** Mohammed A. Dakhil, Gökhan Aydınlı, Johannes Helder, Sevilhan Mennan, Saša Širca, Barbara Gerič Stare, Misghina Goitom Teklu, Daniel P. Bebber

**Author notes:** CONTACT INFORMATION: Daniel P. Bebber, Department of Biosciences, University of Exeter, Exeter EX4 4QD, UK.

## Abstract

Root-knot nematodes (RKN; *Meloidogyne* spp.) are among the most damaging plant-parasitic nematodes worldwide and present a growing threat to global food production as their geographical ranges expand. Despite their importance, the geographical distributions, proliferation risks and crop-level impacts of major RKN species remain poorly understood. Here we assess global habitat suitability for six economically significant or regulated ‘tropical’ RKN species: *M. arenaria*, *M. incognita*, *M. javanica*, *M. enterolobii*, *M. ethiopica* and *M. luci*. We integrated statistical species distribution models (SDMs), mechanistic thermal-time development models and experimentally derived host-suitability metrics. Ensemble SDMs, parameterised using global occurrence records and soil-derived bioclimatic variables, indicate broad climatically suitable ranges for the four well-represented species (*M. arenaria, M. incognita*, *M. javanica* and *M. enterolobii*) across much of the tropics, subtropics and warm temperate regions. Suitability was lower in cooler northern latitudes and, to a degree, in the humid tropics. Predictions for *M. ethiopica* and *M. luci* remain uncertain due to limited occurrence data. Thermal-time phenological models, built from published estimates of base temperatures and Growing Degree-Day (GDD) requirements, revealed a strong latitudinal gradient in potential generation number, with life-cycle completion unlikely at higher latitudes. The risk to 26 globally important crops was evaluated by combining crop-specific reproduction factor and gall index data with global crop distribution maps, indicating highest potential impacts in southern Brazil, the central United States, parts of West and East Africa, eastern India and northern China. Together, our analyses show that most global agricultural land is suitable for the focal species’ establishment and proliferation, with risk shaped jointly by climate, soil conditions and crop hosts. Strengthening distributional surveys and experimentally quantifying species-specific thermal and host responses will be essential to anticipate and mitigate future threats from these crop pests under climate change.

## INTRODUCTION

Crop pests and diseases which threaten food security are shifting their distributions and impacts in response to global change (Ma et al., 2025; Singh et al., 2023). Observations show poleward range expansions of crop pests and diseases in recent decades (Bebber et al., 2013), with further spread likely (Chaloner et al., 2021; Pequeno et al., 2024). Insect pest damage will rise as warming accelerates their metabolism and growth (Deutsch et al., 2018). Plant parasitic nematodes (PPN) that feed on plant roots are among the most damaging crop pests worldwide (Lilley et al., 2024).

Nematodes (Animalia, Phylum Nematoda), commonly known as roundworms, are a diverse and abundant group of parasitic and free-living animals found in nearly all ecosystems, from oceanic trenches to desert soils. Among soil-borne nematode ecological guilds, PPN are thought to be second-most abundant after bacterivores (van den Hoogen et al., 2019) and over four thousand species have been described (Decraemer & Hunt, 2024). Among natural ecosystems they tend to be most abundant in temperate broadleaf forests (van den Hoogen et al., 2019), but their distributions in managed, i.e. agricultural, ecosystems remain largely unexplored. A major challenge in assessing crop losses to PPN has been the difficulty of detecting and identifying the species responsible, especially in regions with limited phytosanitary capacity (Coyne et al., 2018). Even when nematode damage is suspected, species identification based on morphology is challenging in most cases and requires substantial technical expertise (Coyne et al., 2018). Hence, a better understanding of PPN identification, distributions, impacts, and responses to environmental change is an important goal in ensuring future food security.

Root-knot nematodes (RKN) in genus *Meloidogyne* are recognized as the most damaging group of PPN (Jones et al., 2013). Root gall (‘knot’) formation in the host is induced by second stage juveniles which inject effector compounds into protoxylem cells to form multinucleate ‘giant cells’ upon which they feed (Rutter et al., 2022). Their very wide host range, parthenogenetic reproduction, the difficulty in breeding resistance in crop species, and challenges in utilization of chemical controls for soil-borne pests, makes RKN among the most feared threats to plant health (Jones et al., 2013; Trudgill, 1997).

Hazardous nematicides such as aldicarb and oxamyl are increasingly prohibited, and growers are prioritizing health and safety when selecting control methods (Talavera et al., 2024). Given the limitations of chemical controls, cultural methods, such as raising soil temperatures to around 40°C through solarization, and microbial biocontrols, are being explored (Vashisth et al., 2024). Unlike many insect pests and fungal diseases of plants, PPN do not have an aerial dispersal stage but are disseminated largely through trade in plant products (Banks et al., 2018). Thus, quarantine and phytosanitary actions remain the most valuable defence against the further spread of damaging PPN, both within and across national borders.

More than 100 species of RKN have been described (Hodda, 2022), with *M. luci* being the most recently reported (Bačić et al., 2023; Gerič Stare et al., 2017). The European and Mediterranean Plant Protection Organization (EPPO) A1 list of pests currently observed only outside the EPPO region contains six PPN, including *M. ethiopica* (but see Felek & Akyazi, 2024). The EPPO A2 list of pests known to have a limited distribution within the EPPO region contains 14 PPN, including *M. chitwoodi*, *M. enterolobii*, *M. fallax*, *M. graminicola*, *M. luci* and *M. mali*. In addition, three further ‘tropical RKN’ species (sensu Álvarez-Ortega et al., 2019), *M. incognita*, *M. arenaria* and *M. javanica*, appear to be expanding their ranges rapidly into Europe from established populations in the Mediterranean basin (Bebber et al., 2014; Rusinque et al., 2023). Here we focus on six species within Clade 1 ‘tropical RKN’ (Álvarez-Ortega et al., 2019): *M. incognita*, *M. arenaria* and *M. javanica* within the Incognita Group; and *M. enterolobii*, *M. ethiopica* and *M. luci*. Hereafter, these species are collectively termed Tropical RKN (TRKN).

Protected cultivation settings, e.g. glasshouses where warm temperatures are maintained, are likely to have facilitated TRKN establishment in Europe (Talavera et al., 2012). Temperature is a major determinant of RKN life-cycles, with TRKN thriving under warmer conditions (Subbotin et al., 2021; Vashisth et al., 2024). Accordingly, RKN phenology can be described using thermal time models, in which daily temperatures above a baseline contribute to growing degree days (GDD) and accumulation of a certain threshold number of GDD predicts completion of a generation (Trudgill & Perry, 1994). RKN may be killed by temperatures below some minimum, which in thermophilic TRKN may be as high as 10°C (Subbotin et al., 2021). Thus, warming due to climate change may exacerbate crop losses by enabling invasion of hitherto unsuitable regions, increasing the number of TRKN generations per growing season, and enhancing overwintering survival probabilities over a larger geographical range.

Many crop pests and pathogens appear to have expanded their ranges in response to climate change (Bebber, 2015). However, global disparities in surveying and identification capacity may have obscured these trends. On average, recorded PPN occurrences have shifted toward the equator, possibly reflecting earlier detection in wealthier, higher-latitude countries (Bebber et al., 2013).

Moreover, species range responses to climate change can follow complex, non-linear patterns (Lehmann et al., 2020; Rubenstein et al., 2023). Despite their agricultural significance, the likely impacts of climate change on PPN distributions remain poorly understood. The emergence of *M. luci* and *M. incognita* in Europe suggests that warming may have enabled their establishment in previously unsuitable areas. Here, we evaluate both the realised and potential climatic suitability of Europe for six TRKN species using two complementary modelling approaches: (i) species distribution models (SDMs), which use statistical associations between known occurrences and environmental conditions, and (ii) mechanistic thermal models, based on temperature-dependent development and survival thresholds.

## METHODS

### TRKN species

We analysed the distributions of six TRKN species: three ‘major species’ (Elling, 2013; Moens et al., 2009) with high economic impact within the Incognita Group clade (Álvarez-Ortega et al., 2019), *M. incognita, M. arenaria* and *M. javanica*; and three species subject to plant-health quarantine or listed by the European and Mediterranean Plant Protection Organization (EPPO) because of their recognised or potential biosecurity risk (*M. enterolobii*, *M. luci* and *M. ethiopica*). *M. arenaria* is considered to have global distribution in regions where the average temperature of the warmest month is 36°C or below and the average temperature of the coolest month is 3°C or above (CABI, 2021). It is considered rare where the average annual temperature is below 15°C, and most common where average annual precipitation is 1000–2000 mm. *M. incognita* is thought to have a global distribution and is endemic across much the tropics and subtropics (Eisenback, 2020). *M. javanica* is another warm climate species which has been largely restricted to greenhouses in Europe. It appears to be more common in drier areas (CABI, 2022). *M. enterolobii* is thought to have a more limited known distribution in warmer regions than the preceding species, and is thought not to be able to survive outside protected cultivation in colder climates (Castillo & Castagnone-Sereno, 2020; Pan et al., 2023). *M. ethiopica* is thought to be restricted to Africa, South America and Türkiye, and appears to be able to survive in temperate conditions (Carneiro & Carneiro, 2012; Felek & Akyazi, 2024). *M. luci* is a recently described species with a widespread distribution (Supplementary Table S1). *M. ethiopica* isolates in Europe have been reclassified as *M. luci* (Gerič Stare et al., 2017).

### Implications of TRKN evolution for distribution modelling

All six TRKN reproduce by obligate mitotic parthenogenesis and are allopolyploid (Dai et al., 2023). Before considering their global distributions, we note that species identification within TRKN is challenging due to their reticulate evolutionary histories involving multiple hybridization events and very low divergence of mitochondrial genomes (Dai et al., 2023). These features are associated with very similar morphologies and broadly overlapping phenotypes among species, and have implications for the reliability of species-level occurrence records. Consequently, some records may be misidentified (for example, confusion between *M. ethiopica* and *M. luci*). Thus, while we model the distributions of each species separately, we prioritised consistency of environmental predictors and modelling structure across species to facilitate direct comparison and synthesis, rather than species-specific optimisation, and subsequently combine individual risk maps to provide an overall estimate of environmental suitability for the six species as a group.

### Distribution data

We obtained occurrence data of the target six *Meloidogyne* species from seven sources: the Global Biodiversity Information Facility (GBIF; https://www.gbif.org), CABI (https://www.cabidigitallibrary.org/), EPPO (https://gd.eppo.int/), NCBI (https://www.ncbi.nlm.nih.gov/), and relevant literature (peer-reviewed journal articles, dissertations and reports). Locations from distribution map images were abstracted using Google Earth Pro software (https://www.google.com/intl/en_uk/earth/about/versions/#earth-pro). Only point locations were included, i.e. we did not include national or sub-national region data. Location data were cleaned by removing duplicates, points located in oceans, and points located in non-cropping areas based on global crop distribution (Potapov et al., 2022), first checking if there had been a possible switch between latitude and longitude for apparently erroneous locations.

### Data handling, analysis and visualization

All analyses were performed in the R environment v.4.5.0 (R Core Team, 2025). Manipulation of spatial data was undertaken using the *terra* package v.1.8–60 (Hijmans et al., 2025) for R, with *tidyverse* v.2.0.0 (Wickham et al., 2019) for additional packages facilitating data manipulation.

Figures were created using the *ggplot2* package v.3.5.2 (Wickham et al., 2024) and *cowplot* v.1.2.0 (Wilke, 2025), with *tidyterra* v.0.7.2 (Hernangómez, 2023) for spatial raster and vector data and *biscale* v.1.1.0 (Prener et al., 2025) for bivariate maps. Land mass and country outlines were obtained from the GADM database (https://www.gadm.org) using the *geodata* package 0.6.2 (Hijmans et al., 2024). We developed ensemble species distribution models (SDMs) using the *sdm* package v4.4.1 (Naimi & Araújo, 2016). The *sdm* package employs the following R packages to implemented the models we tested: Package *stats* for Generalized Linear Models (GLM); *mgcv* (Wood, 2017) for Generalized Additive Models (GAM); *gbm* (Ridgeway et al., 2024) for Boosted Regression Trees (BRT); *kernlab* (Karatzoglou et al., 2004) for Support Vector Machines (SVM); *randomForest* (Breiman et al., 2024) for Random Forests; and *earth* (Milborrow et al., 2024) for Multiple Adaptive Regression Splines (MARS). We used the Javascript implementation of the MaxEnt algorithm (Phillips et al., 2006), available from https://biodiversityinformatics.amnh.org/open_source/maxent/. For visualization, global maps were reprojected to the Equal Earth area-preserving coordinate reference system (Šavrič et al., 2019) and resampled using the mean to a 20 km or 40 km grid, corresponding to approximately 300 dpi print resolution. This aggregation reduced visual sparsity caused by patchy cropland at the original 3 km resolution. European maps were restricted to longitude 10° W–33° E and latitude 33° N–72° N, then reprojected to the area-preserving ETRS89 / LAEA Europe coordinate reference system (EPSG:3035) at 4 km resolution. These maps are intended to illustrate broad geographical patterns rather than fine-scale suitability at field resolution.

### Environmental variables

Previous studies have identified temperature, precipitation, and soil properties as potentially important correlates of global TRKN distributions (Sasser et al., 1983) and soil nematode abundance (van den Hoogen et al., 2019). We employed soil (rather than air) temperature-derived variables as predictors. Mean monthly soil temperature estimates and 11 soil temperature bioclimatic variables (SBIO1–11) at 30 arc-seconds spatial resolution and two depths (0–5 cm and 5–15 cm) for the period 1979–2013 were obtained from the Global Maps of Soil Temperature database (Lembrechts et al., 2022). We detected calculation irregularities in the isothermality dataset (SBIO3), so this variable was omitted from further analysis. Variable values at the two depths were highly correlated (Pearson correlation > 0.90 for most variables, 0.82 for mean diurnal range), so we used the layer thickness-weighted mean for modelling. Precipitation bioclimatic variables (BIO12–BIO19) at 30 arc second resolution for the period 1981–2010 were obtained from CHELSA v2.1 (Karger et al., 2017, 2018). Although the Global Maps of Soil Temperature variables were derived from CHELSA v1.2 and precipitation from v2.1, this version mismatch is unlikely to materially affect our results, as broad-scale precipitation seasonality patterns are consistent between versions and SDMs are inherently correlative. We obtained 9 soil textural and chemical variables from the International Soil Reference and Information Centre (ISRIC) SoilGrids250m, aggregated to ∼1 km resolution (Hengl et al., 2017). We resampled (mean values) all environmental variable datasets to 0.025° (approximately 3 km^2^) resolution to match the Global Land Analysis & Discovery (GLAD) global overview cropland distribution (Potapov et al., 2022). TRKN are almost exclusively sampled from agricultural crops, therefore we omitted non-cropland regions from further analysis. Predictor variables data sources are summarized in Supplementary Table S2.

### Ensemble species distribution modelling and model accuracy

For each species, 5000 background points were randomly sampled from cropland areas within 1000 km of known presence locations to approximate the species’ accessible area (Barve et al., 2011) and help reduce sampling bias (Barbet-Massin et al., 2012; Phillips et al., 2009). Environmental predictor values were extracted from the presence and background points. Multicollinearity among predictors was avoided by stepwise removal of variables with Variance Inflation Factor (VIF) greater than 5 (Guisan et al., 2017), with the exception of mean annual temperature (SBIO1) and mean annual precipitation (BIO12) which we retained as representing major climatic gradients. Predictor selection by VIF was conducted on the combined presence and background data for all six TRKN species, and the same predictors were used to model each species’ distribution. This approach was adopted to ensure comparability of response curves, suitability patterns and derived richness estimates across species. Model performance was first evaluated by five-fold cross-validation using seven species distribution modelling (SDM) algorithms. The mean True Skill Statistic (TSS) and Area Under the receiver operating characteristic Curve (AUC) were compared across algorithms, and the three best-performing methods across all species were selected for 30-fold bootstrap resampling, using 70 % of records for training and 30 % for validation in each iteration. The same set of algorithms was retained for all species to ensure that differences in predicted suitability reflected species–environment relationships rather than differences in modelling structure, while species-specific model performance was accounted for through TSS-weighted ensemble predictions.

Final models were fitted using all presence and background data. Ensemble suitability predictions for each species were obtained by weighting the predicted suitabilities of the three algorithms by their bootstrap mean TSS. Predictor-suitability response curves were calculated from the bootstrap means for each algorithm, and final ensemble response curves were derived as TSS-weighted means across algorithms. These curves are presented for qualitative comparison of trends only, as absolute response magnitudes are algorithm-specific. The importance of each variable in each bootstrap replicate was calculated as the reduction in AUC when the values of that variable were randomly permuted while keeping all other predictors unchanged. Binary suitability predictions were obtained using two habitat suitability thresholds: the Minimum Training Presence (MTP) threshold, defined as the lowest suitability value associated with any training presence point; and the tenth percentile training presence (P10) threshold which excludes regions with suitability values lower than those of the lowest ten percent of occurrence records (Pearson et al., 2007). The MTP threshold was selected for final suitability mapping because several TRKN species are known to occur widely (Luc et al., 1990), but verified georeferenced records remain sparse. In such cases, more restrictive thresholds such as P10 risk excluding environmentally suitable but under-sampled regions (Pearson et al., 2007).

### Thermal time phenological models

Thermal time models, also known as heat sum or accumulation models, are commonly used to predict phenology in a range of organisms including nematodes (Trudgill, 1995; Trudgill & Perry, 1994; Velloso et al., 2022) and their crop hosts (Zhao et al., 2019). Thermal time is expressed as growing degree-days (GDD), the daily accumulation of degrees Celsius above a minimum baseline temperature *T_b_* required for development, with phenological stages reached once species-specific GDD thresholds are met. GDD is normally calculated from the average of daily minimum and maximum temperatures (McMaster & Wilhelm, 1997). In this case, we calculated observed GDD sum (*S_o_*) from mean monthly soil temperatures *T_m_* at each soil depth using the simplification that each day of the *i*th month has the mean monthly temperature *T_m,i_* :

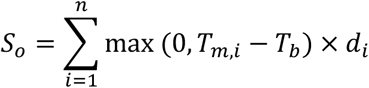

where *n* is the duration of the period being considered (usually 12 for the entire year, but can be restricted to particular host plant growing seasons), *d_i_* is the number of days in the *i*th month, and max indicates that when *T_m,i_* is below *T_b_* no GDD are accumulated. The *ENVIREM* package for R (Title & Bemmels, 2018) contains a function for this calculation, but the implementation is slow so we wrote a new, faster function avoiding loops. The number of possible generations per year *G* for each was estimated by:

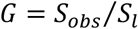

where *S_l_* is the GDD sum required to complete the lifecycle.

Experimentally-derived *T_b_* and *S_l_* estimates for target TRKN species were collated from the literature. Studies which did not measure development over the full life cycle (usually considered as J2 to J2 stage) were omitted (e.g. Dávila-Negrón & Dickson, 2013; Tzortzakakis & Trudgill, 2005). One study found that development slowed above 30°C (Madulu & Trudgill, 1994). However, given the rarity of such high soil temperatures in cropland regions, we did not adjust the model to account for this phenomenon.

As the focus of our study is crop production, we restricted calculations to global cropland-containing grid cells (Potapov et al., 2022). In many crop models, *T_b_* varies among crop species (and sometimes cultivars) and is greater for tropical crops like cassava and banana than temperate crops like wheat and potato (FAO & IIASA, 2022 Appendix 4-3; Zhao et al., 2019). TRKN have high *T_b_* (> 10 °C), similar to tropical crops and greater than temperate crops, for example potato with *T_b_* = 4 °C (Zhao et al., 2019). This implies that TRKN will be active within the growing seasons of most crops. However, crops may be planted later and harvested earlier than expected solely from sufficiently warm conditions (Bondeau et al., 2007; Sacks et al., 2010). Therefore we estimated *G* for a major tropical crop (maize) and temperate crop (potato) based on MIRCA-OS crop calendars for the year 2015 (Kebede et al., 2025). While TRKN have very wide host ranges, maize and potato are widely planted and are vulnerable to these nematodes (Coyne et al., 2018). Crop-specific thermal time accumulation was restricted to months in which each crop (either irrigated or rainfed) was present in each grid cell.

### Crop suitability

The suitability of particular plants as hosts varies among and within TRKN species (Luc et al., 1990). For example, tomatoes are considered to be highly suitable hosts for several of our focal species, but resistant varieties are available (Verdejo-Lucas et al., 2013). To provide a general overview of geographical patterns of crop host suitability, we gathered data on the reproduction factor (RF) and galling index (GI) for 26 of the most common crops globally, for which such information was available (Supplementary Data S1). Only data from non-resistant varieties was used, and where multiple results per crop were reported per study, the mean was taken. RF is the ratio of the final to initial nematode population (RF = *P_f_*/*P_i_*), and GI is a semi-quantitative measure of galling severity (e.g. Maleita et al., 2022). GI is often recorded on a scale of 0–5, but may use other scales (e.g. 0–10) and if so, values were rescaled to the 0–5 range. Crop suitability was mapped using the SPAM crop distribution estimates (IFPRI, 2024), by calculating an area-weighted crop suitability index *C* for both RF and GI per grid cell:

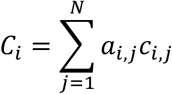

where *a* is the fractional area of the *j*th crop in the *i*th grid cell, and *c* is the mean (across focal TRKN species) of the nematode multiplication metric (RF or GI) values.

Host suitability data were not available for all crop × TRKN species combinations. The index *C_i_* therefore represents the aggregate crop suitability index of host-mediated vulnerability to focal TRKN as a group. This approach is appropriate given the broad host ranges shared by TRKN species and allows consistent, landscape-scale comparison of host availability. This ensemble-based index is intended to capture the vulnerability of cropping systems to TRKN pressure as a whole, rather than the performance of any single TRKN species on a given crop, and therefore remains informative even where individual species show contrasting virulence patterns across hosts.

## RESULTS

### Species distribution models

Precise locations (i.e., exact latitude and longitude) were available for *M. arenaria* (150 observations)*, M. incognita* (326), *M. javanica* (317) and *M. enterolobii* (152) (Supplementary Figure S1, Supplementary Data S1). Only 31 precise locations were available for *M. ethiopica*, in South America and Türkiye, while 45 were available for *M. luci*, in Europe, Türkiye, Brazil and Ethiopia. VIF-based selection retained 15 environmental predictors for SDM, including four soil temperature-derived variables, five precipitation variables, and six physical and chemical variables (Supplementary Table S3). Mean annual soil temperature (SBIO1) and precipitation (BIO12) had slightly higher VIF than other predictors we retained. Five-fold cross-validation of SDMs revealed little variation in model performance among the seven candidate algorithms (Supplementary Table S4). The best algorithm was RF, but inspection of mean response curves of suitability by different predictors revealed highly erratic behaviour indicative of overfitting, and unlikely to approximate species’ responses to the environment. Hence, we selected the three next best algorithms by TSS (MaxEnt, SVM and MARS) for ensemble model fitting.

The three algorithms had similar bootstrap mean TSS scores for each species, hence contributing approximately equally to the final SDM habitat suitability predictions (Supplementary Tables S5, S6). However, the predictor variable importance values differed substantially among algorithms within species (Supplementary Figure S2). SBIO1 was the most important variable or *M. arenaria* and *M. incognita* in all three algorithms, and among the most important predictors for *M. javanica* and *M. luci*. There was no clear or consistent difference in the importance of soil temperature, precipitation or physicochemical predictors, though precipitation was somewhat more important for *M. ethiopica* and climatic variables somewhat more important than physicochemical properties for *M. javanica* and *M. incognita*. Similarly, suitability response curves per predictor varied among models and species, though with some areas of agreement (Supplementary Figure S3). For *M. arenaria*, the three algorithms showed similar responses for SBIO1 (peak ∼ 15–20 °C) and BIO12 (peak ∼ 750–1500 mm), but qualitatively different responses for other predictors such as SBIO4 (soil temperature seasonality) and BIO15 (precipitation seasonality). For *M. enterolobii*, all three algorithms showed declining suitability with BIO12 and SBIO2 and a peak response at around 300–500 mm for BIO19. For *M. ethiopica*, suitability increased with AWC, BIO14 and BIO15 for MaxEnt and MARS. For *M. incognita*, the three algorithms agreed most strongly for BIO12 (peak ∼ 750 mm), BIO15 (increasing), BIO19 (peak ∼ 500 mm), CEC (peak ∼ 20 cmol(+) kg^−1^), SBIO1 (peak ∼ 25–30 °C) and SBIO8 (declining). For *M. javanica*, the algorithms showed similar responses for BIO12, and increasing response with BIO15 and SBIO4, a peak SBIO1 of around 25–30 °C and peak SBIO2 of around 3–5 °C. There was little agreement among models for *M. luci* distributions.

Ensemble model habitat suitability predictions (Supplementary Figure S4) were converted to binary maps using suitability thresholds (Supplementary Table S7) to delineate regions likely to be suitable or unsuitable for each species (Figure 1). Taking the MTP threshold, suitable areas for *M. arenaria*, *M. enterolobii*, *M. incognita* and *M. javanica* spanned parts of the USA, much of Brazil, West and East Africa, southern Europe, China, India and Australia. The humid tropics (the Amazon basin, Congo Basin and Southeast Asia) were less suitable for *M. arenaria* and *M. incognita*. The P10 threshold was far more restrictive, particularly for *M. arenaria*. The predicted suitable areas for *M. ethiopica* and *M. luci* were much smaller, but included parts of the USA, Europe, Brazil, East and West Africa, China and Australia. In general, northern USA and Canada, northern and central Europe, central Asia and Russia, and parts of Indonesia were predicted to be less suitable for TRKN. *M. enterolobii* and *M. incognita* had suitable areas in northern Europe and the British Isles. Overlaying the MTP-thresholded suitability maps for the six species provided a general indication of TRKN habitat suitability. Regions determined to be suitable for the greatest number of TRKN were southeastern USA, southern Brazil, southern Europe and Türkiye, West and East Africa, eastern China and southern Australia (Figure 2). Within Europe, greater suitability was predicted in warmer, wetter regions with warmer winters, including the Atlantic coasts of Spain, France and the British Isles, as well Mediterranean regions and Türkiye (Supplementary Figure S5).

**Figure 1.**
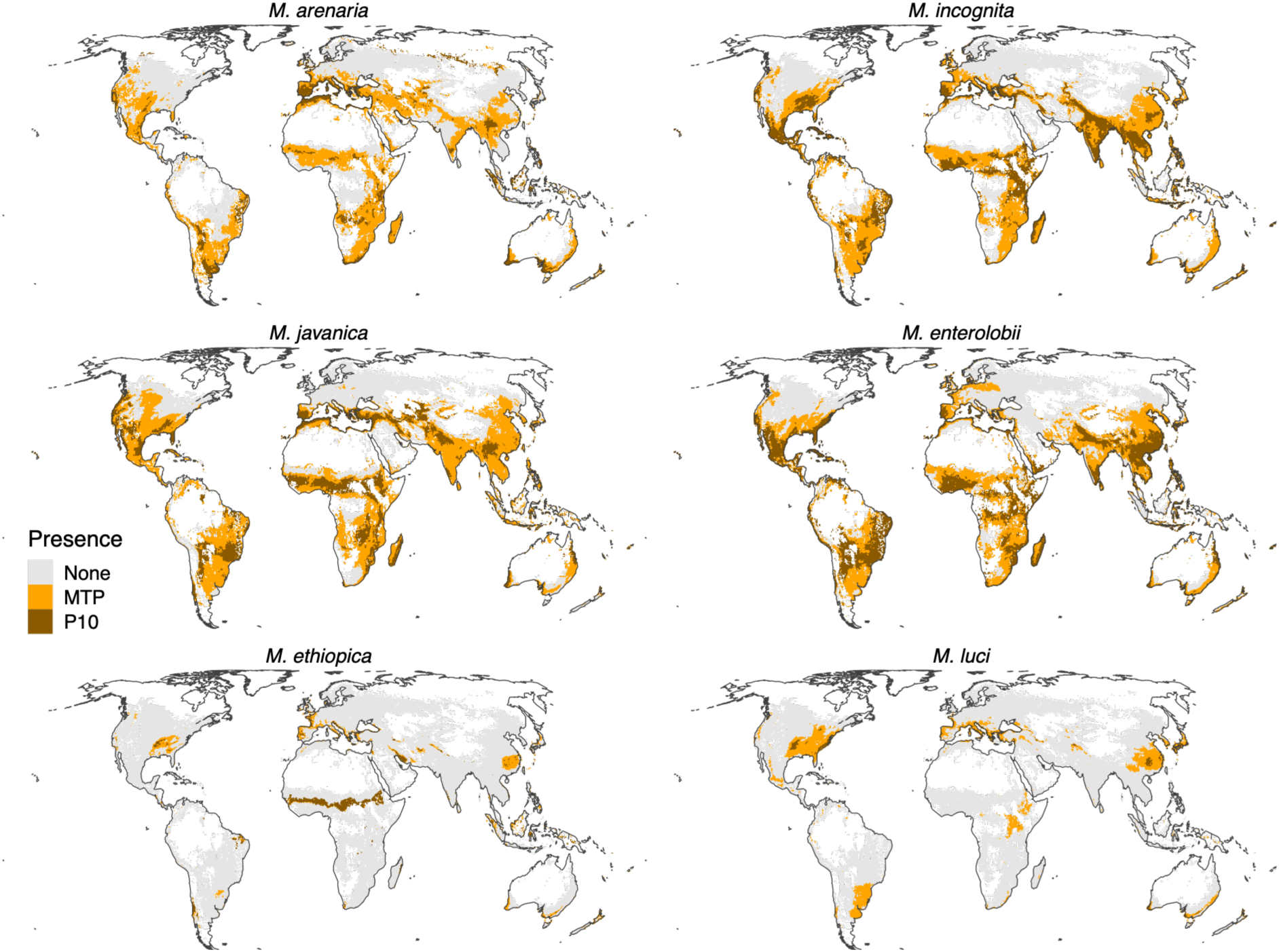
Binary habitat suitability by species. Minimum Training Presence (MTP; orange) and 10th-percentile training presence (P10; brown) thresholds were estimated from occurrence records and ensemble suitability predictions.

**Figure 2.**
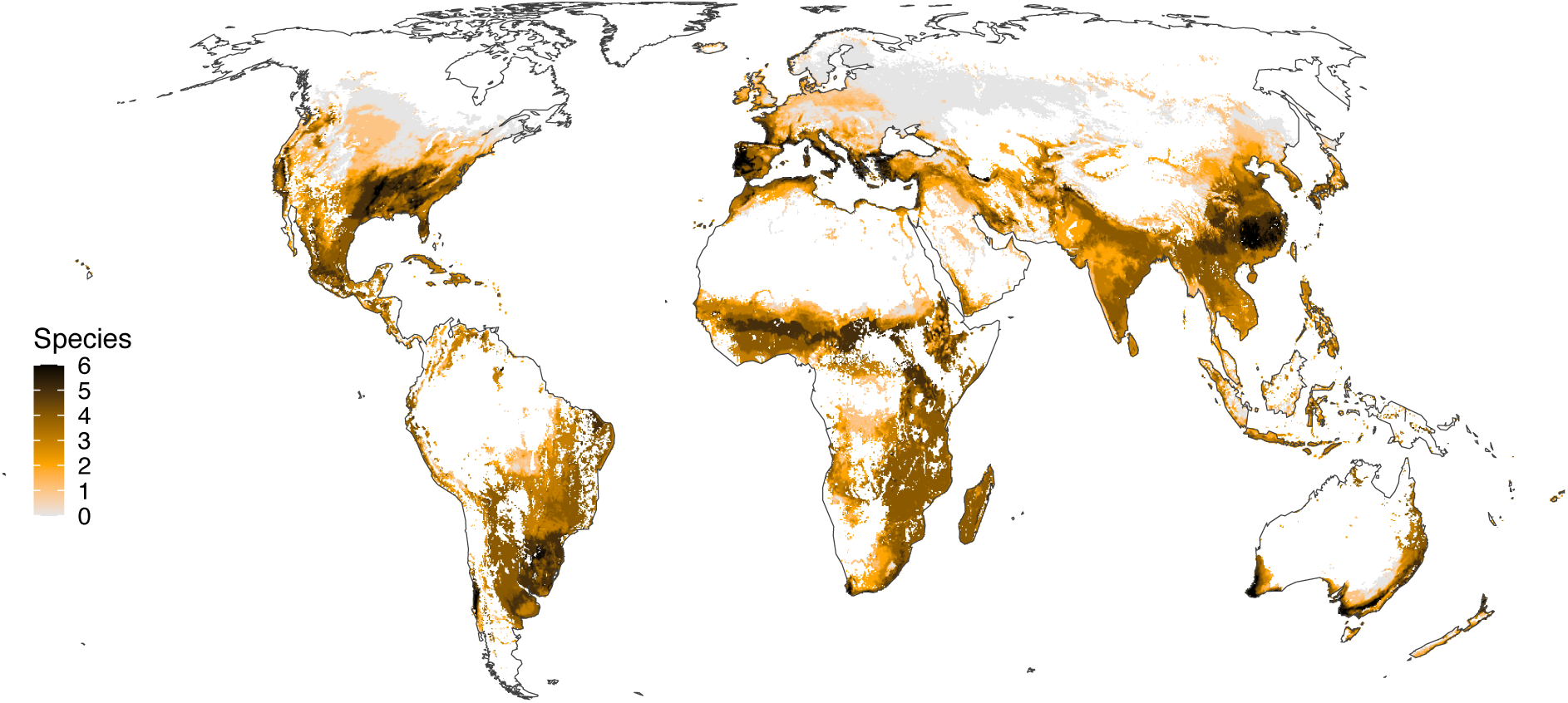
Combined suitability for six TRKN species. Colours indicate the number of species predicted to occur under the MTP threshold (grey indicating zero).

### Phenological models

Life cycle GDD sums (*S_l_*) were available for three target TRKN species (*M. enterolobii, M. incognita* and *M. javanica*) and varied between 350 and 552 °C.day with base temperature (*T_b_*) between 10.0 and 17.2 °C (Supplementary Table S8, Supplementary Figure S6). In comparison, two PCN species (*G. pallida* and *G. rostochiensis*) had a similar *S_l_* range (398–579 °C.day) but substantially lower *T_b_* (4.0–6.0 °C). In most cases, *S_l_* and *T_b_* had been estimated by regressing the reciprocal of life cycle duration (in days^−1^) against temperature, under experimental conditions with constant temperature and a variety of hosts. In most experiments, only a small number of temperatures (around four settings) had been employed with no replications per temperature, and no uncertainty estimates are provided for the parameter estimates. The majority of TRKN *S_l_* values fell around 475 °C.day with *T_b_* around 11.0 °C. Two estimates for *M. javanica* showed higher *T_b_* and lower *S_l_* than other studies (López-Gómez et al., 2014), while one estimate for *M. incognita* had a lower *S_l_* (Ploeg & Maris, 1999). Insufficient replicates within species were available for statistical analyses of variation among species, or the effects of nematode geographical origin or host plant species.

The mean number of potential annual TRKN generations in global croplands, averaged over the 11 estimates from the literature, varied from zero to over 20 (Supplementary Figure S7). There was no indication of substantial variation in generation times among TRKN species or crops, for the 11 combinations found in the literature. The number of potential annual generations varied slightly among TRKN species, host plant species and study. No study compared phenology of a particular TRKN species on different hosts, so direct analysis of the effect of host on development rate was not possible. Slightly fewer generations were predicted for both *M. incognita* and *M. javanica* on *Cucurbita melo* than on other hosts. However, the difference was of a similar order of magnitude to the difference between two studies conducted on *M. incognita* with *S. lycopersicon*.

Slightly more generations were estimated at 0–5 cm soil depth than 5–15cm (median difference 0.5 months), though with substantial spatial variability (Supplementary Figure S8). More generations were estimated from mean monthly air temperatures than from 0–5 cm soil depth temperatures in parts of the USA, Brazil, central and East Africa, China and Southeast Asia (Supplementary Figure S8). Fewer generations were estimated using air than soil temperatures in the Sahel, the Middle East, parts of India and Australia. There was a very strong latitudinal gradient in mean annual generations, with the highest values predicted in West Africa and the Sahel, the Middle East (Syria, Iraq and Iran), India and Southeast Asia and the Philippines (Supplementary Figure S9a). Only Canada, northern Europe and the Russian Federation contained substantial areas of cropland where the mean number of generations per year was below one, i.e. no life cycle completion. The variation among model estimates increased with the mean (Supplementary Figure S9b).

Restricting phenological models to crop calendars reduced the potential number of generations, though the overall latitudinal trends remained (Figure 3). For the maize crop calendar, completion of at least one life cycle (mean of 11 models) within the host growing season was possible everywhere except in parts of Canada and the US, the Andes, north Africa, Portugal, northern France and the UK, most of northern Europe and Russia, parts of Ethiopia, small areas of India and much of China. For the potato crop calendar, life cycle completion was not predicted to occur in areas of Canada, the Andes, northern Europe and Russia, parts of Ethiopia, Afghanistan, western China and southern Australia. In Europe, most of Spain, France, Italy, and the Balkans were estimated to be suitable for completion of at least one generation during the potato cropping season (Figure 4). Minimum soil temperatures (BIO6) remain above freezing across most of western Europe at both 0–5 cm and 5–15 cm, whereas eastern Europe and mountainous regions reach freezing point and below (Supplementary Figure S10).

**Figure 3.**
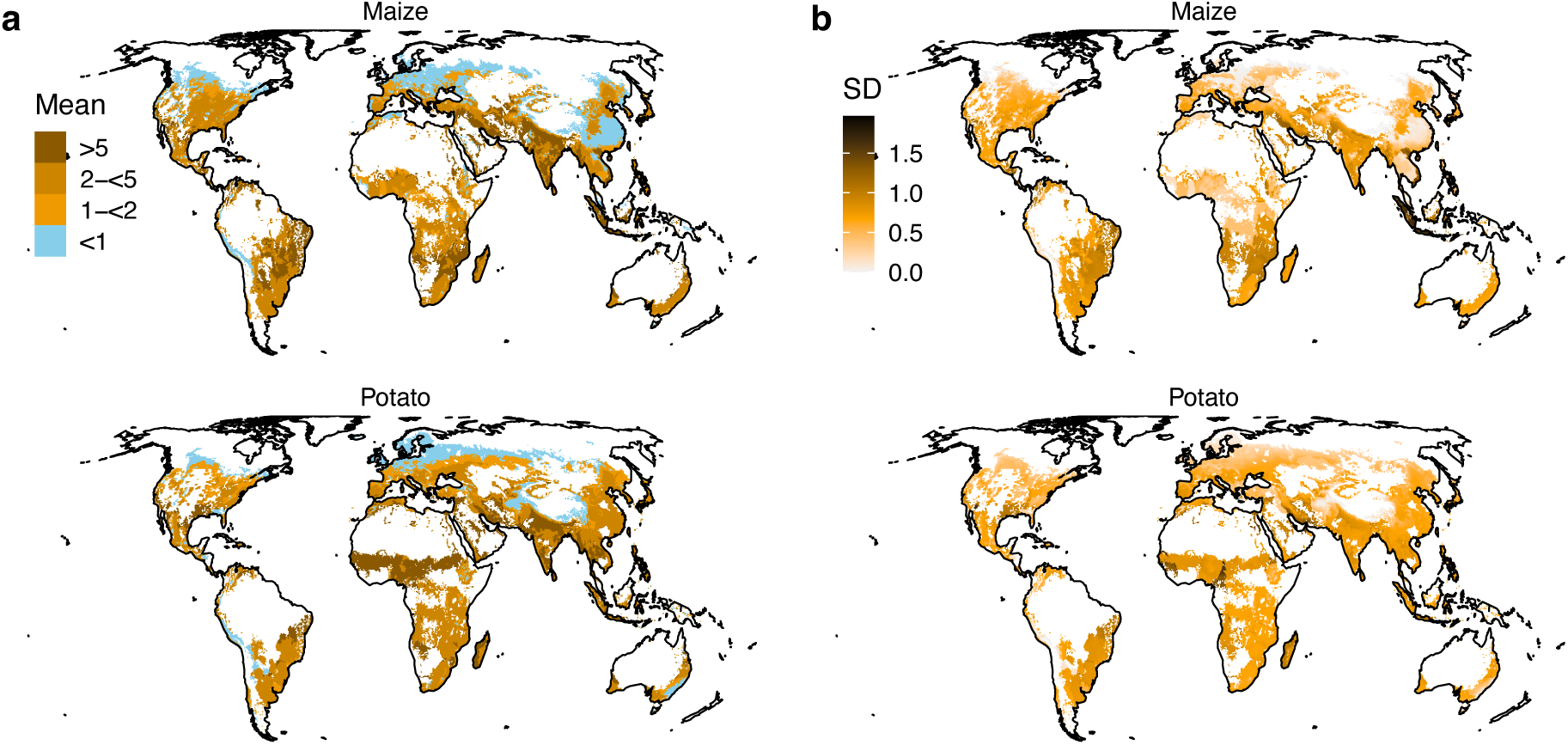
TRKN generations by maize and potato crop calendar. a) Mean life cycles per year estimated from mean monthly soil temperatures at 0–5 cm during maize and potato growing seasons. b) Standard deviations of mean life cycles per year.

**Figure 4.**
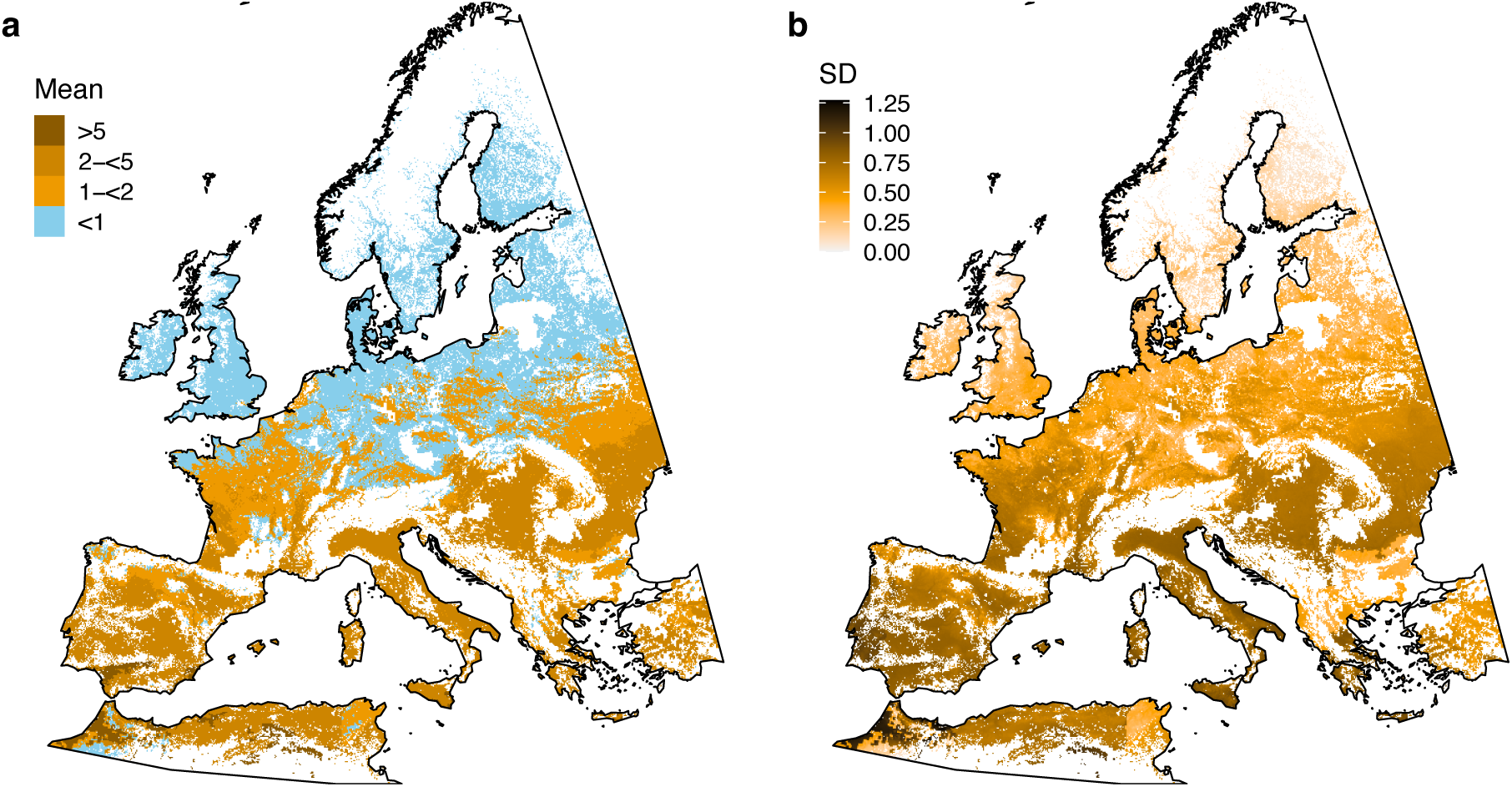
TRKN generations on potato in Europe. a) Mean generations during the potato growing season. B) Standard deviation among model estimates. Estimates are for all potato grid cells in the MIRCA-OS crop calendar dataset.

### Crop suitability

We found experimentally-derived estimates of RF and GI for 26 of the 42 crops analysed in the SPAM global crop distribution maps, together representing 76 % of global cropping area in 2020 (FAO, 2025). Host suitability estimates were available for all focal TRKN species, but not for each crop-RKN combination (Supplementary Data S2). Accordingly, results were pooled to give an ensemble measure of crop suitability for focal TRKN considered as a group. Mean RF per crop varied from below 1 in citrus (*Citrus* spp.), coconut (*Cocos nucifera*), barley (*Hordeum vulgare*), cowpea (*Vigna unguiculata*), yam (*Dioscorea* spp.), cassava (*Manihot esculenta*) and sorghum (*Sorghum bicolor*) to over 30 in bean (various species), Arabica coffee (*Coffea arabica*), tomato (*Solanum lycopersicum*) and potato (*Solanum tuberosum*) (Supplementary Table S9). Mean GI varied from below 1 in citrus, coconut, millet (mainly *Panicum miliaceum*), cowpea and sorghum to over 4 in sweet potato (*Ipomoea batatas*), Arabica coffee, banana (*Musa* spp.) and sugar beet (*Beta vulgaris*).

Mean RF and GI per crop were strongly correlated (Supplementary Figure S11, Spearman’s ρ = 0.76). Globally, crop area-weighted RF and GI were highly correlated (Spearman’s ρ = 0.975). Both metrics were highest in southern Brazil, central USA and Canada, parts of West Africa and East Africa, Romania and Ukraine, western India and northern China (Figure 5a,b). Within Europe, central and eastern regions had generally greater RF and GI (Figure 5c,d).

**Figure 5.**
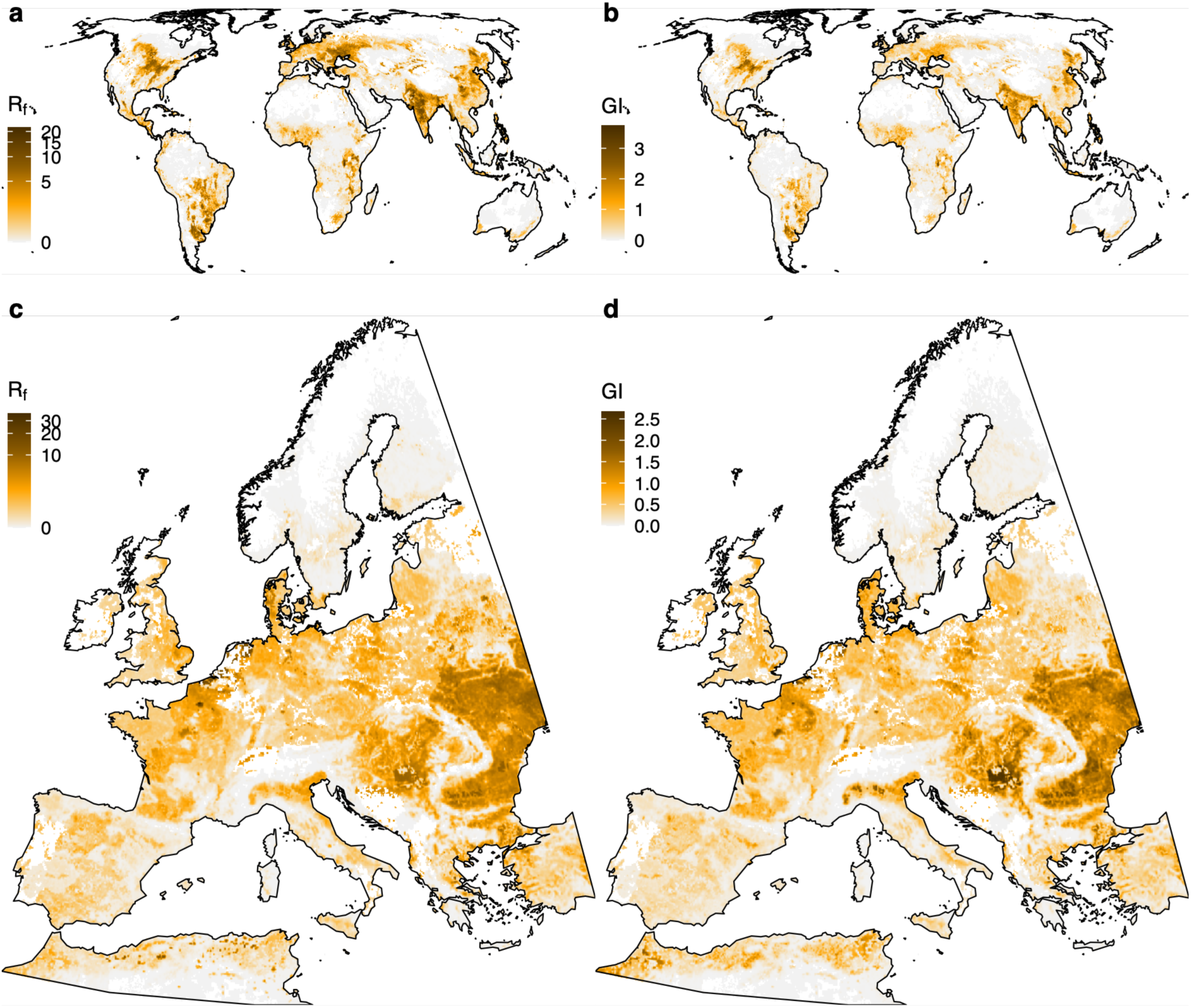
Crop area-weighted TRKN proliferation suitability. Values are means across focal TRKN for 26 crops. a) Reproduction factor (RF), global. b) Galling index (GI), global. c) RF, Europe. d) GI, Europe.

## DISCUSSION

Root-knot nematodes pose a major risk to global crop production, and the geographical ranges of several TRKN species have been expanding in temperate regions. Our analysis combined ensemble SDMs, thermal time phenological models and crop susceptibility estimates to understand where TRKN populations are most likely to establish and proliferate. Our results suggest that the majority of the world’s agricultural lands, excepting cooler northern temperate regions, are likely to be suitable for TRKN establishment. Proliferation risk is then determined by growing season temperatures and the particular crop species under cultivation. Regions with the greatest habitat suitability and crop proliferation risk included Mexico and the Caribbean, southern Brazil, the Mediterranean, East and West Africa, east India and northern China, as indicated by the bivariate synthesis of SDM and crop suitability metrics (Figure 6, Supplementary Figure S12). For species with sufficient distributional information (*M. arenaria*, *M. incognita*, *M. javanica* and *M. enterolobii*), SDMs indicated latitudinally-wide habitat suitability, with cooler temperate croplands and some parts of the humid tropics less likely to support establishment. This agrees with the consensus, for *M. arenaria* at least, of a distribution constrained by climatic extremes (CABI, 2021). Our thermal time generation models confirm a general latitudinal gradient in potential proliferation rates.

**Figure 6.**
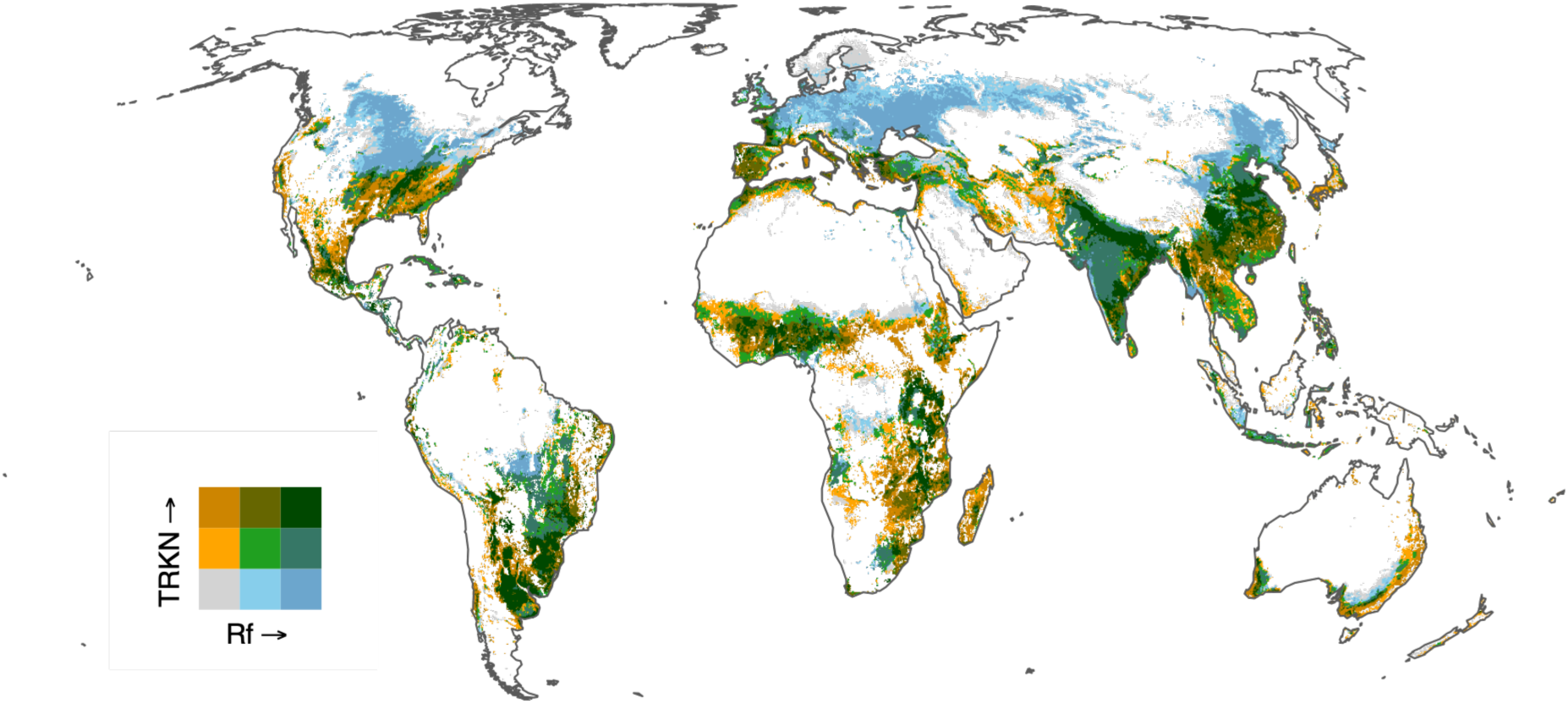
Combined TRKN habitat and crop suitability. Habitat suitability is based on the sum of MTP-threshold SDM predictions, and crop suitability on the crop area-weighted reproductive factor (RF). Colour scales were derived from quantiles of habitat suitability and inverse hyperbolic sine-transformed RF. Blue indicates high habitat but low crop suitability, orange indicates low habitat but high crop suitability, and green indicates regions where both are suitable for TRKN proliferation. Grey denotes areas unsuitable for both.

Constraints such as temperature extremes, moisture regimes and soil properties are likely to explain areas of disagreement between thermal time models and SDMs regarding establishment potential in the humid tropics. Pinpointing the major distributional constraints was not possible from our SDMs, given that temperature, precipitation and soil composition variables all contributed in different ways to modelled distributions. In general, all three factors are correlates of both soil nematode abundance (van den Hoogen et al., 2019) and TRKN distributions (Sasser et al., 1983). Soil temperature has long been known to strongly influence migration, root penetration, proliferation and survival of TRKN (Bird & Wallace, 1965; Daulton & Nusbaum, 1961; Sasser et al., 1983). Optimal temperatures have previously been estimated at around 25–30°C (Bird & Wallace, 1965), with reduced growth and reproduction below 15°C (Vrain et al., 1978) and above 35°C (Daulton & Nusbaum, 1961; Ploeg & Maris, 1999). Thermal time models, predicated on the approximately linear rate of increase in biological rates with temperature between the minimum and optimum temperatures of a process (Trudgill & Perry, 1994), tend to confirm a lower limit of around 10°C for life cycle completion, compared with around 5° C for PCN. Very high soil temperatures during a crop growing season could potentially reduce nematode populations (Oka, 2019), but heat extremes can also reduce crop productivity directly (Heino et al., 2023).

Determining the relative importance of temperature responses in the host plant versus TRKN on crop yields will require further empirical investigation. Analogously, projected shifts in crop yields under future climate change are likely to be mirrored by fungal and oomycete crop pathogen infection risks, potentially offsetting any benefits of warming at higher latitudes (Chaloner et al., 2021). Modelling climate change effects on TRKN distributions was beyond the scope of the present study, but it is likely that warming will bring hitherto unsuitable regions into the TRKN thermal niche. Indeed, the recent detection of *M. ethiopica* in Türkiye (Felek & Akyazi, 2024), hitherto known only from lower latitudes, suggests that climate change may already be facilitating TRKN invasions.

Soil physico-chemical variables have also been reported as influencing TRKN abundance (de Oliveira Costa et al., 2025). However, the relative importance of soil properties varied among SDM algorithms and among species. Although predictors were VIF-screened to reduce strong linear collinearity, soil variables nonetheless retain shared broad-scale spatial structure and non-linear associations with climate and land-use gradients, which VIF does not address. Different SDM algorithms may therefore exploit different predictors to achieve comparable predictive performance and variable importance rankings should be interpreted cautiously. At finer spatial scales, experimental and field studies have demonstrated clear effects of soil properties on TRKN populations (Narasimhamurthy et al., 2022), but such effects are difficult to resolve in correlative global analyses that necessarily average over substantial local heterogeneity.

Climate change is expected, in general, to shift species distributions poleward as populations track preferred thermal regimes and for several classes of crop pests and pathogens, including fungi, oomycetes and insect clades, there is evidence for such shifts (Bebber et al., 2013). However, historical observations of plant pathogenic viruses and nematodes appear to show the opposite trend, moving toward the equator. Bebber et al. (2013) speculated that this is because these taxa are more difficult to detect and identify, and therefore tend to be reported in countries at higher latitudes which historically have possessed greater scientific and technical capacity. Apparent latitudinal patterns may partly reflect historical sampling and identification biases rather than true biogeographic limits. The case of *M. luci*, first described in 2014 (Carneiro et al., 2014), illustrates this phenomenon. Unique esterase profiles (L3, sometimes coded M3) enable *M. luci* to be identified from earlier studies. The majority of confirmed *M. luci* observations have come from sub-tropical and temperate locations in Europe, Türkiye and Brazil, along with a small number of tropical samples from Latin America and Africa (Supplementary Table S1). While the environmental niches of both *M. luci* and *M. ethiopica* remain poorly resolved, due to their wide host ranges we speculate that further sampling may reveal these species to be widely distributed, and will provide a more detailed understanding of TRKN distributions in general.

TRKN have very wide host ranges, but agricultural crops are better hosts (in terms of shorter nematode generation times and greater fecundity) than wild species and weeds, and some crop species are better hosts than others (Trudgill & Blok, 2001). Solanaceous crops (e.g. tomato, potato, aubergine, pepper, tobacco) are considered good TRKN hosts globally (Luc et al., 1990), excluding varieties bred for resistance (e.g. Ehwaeti et al., 1999). Mean RF and GI per crop were strongly correlated across the 26 crops we tested, and this correlation is seen within individual TRKN species when large numbers of hosts are compared (e.g. Lima et al., 2009). Both proliferation metrics have recognised limitations: RF depends on the initial inoculum density used, while GI is a semi-quantitative measure of root damage that may not always reflect nematode reproduction. Accordingly, we interpret RF and GI as complementary indicators of host suitability at broad spatial scales.

Our analyses were constrained by a lack of both accurate distribution records and biological information, such as phenological models, as well as uncertainties around species identification (particularly for *M. ethiopica* and *M. luci*). Most of these species are thought to be more widespread in the tropics than our precise location records indicate, therefore we employed the inclusive MTP threshold to avoid artificially restricting suitable areas. We also combined GDD, RF and GI information across species as a pragmatic approach given the sparse and inconsistent data available for individual taxa. Despite these constraints, our results provide the most comprehensive global assessment to date of the environmental and agronomic constraints shaping TRKN distributions and impacts. By identifying where establishment and proliferation are most likely, we offer a foundation for improved surveillance, targeted management strategies and projections of future climate-driven risks. Strengthening the empirical evidence base through coordinated surveys and experimental work will be essential for safeguarding crop production in the face of an increasingly pervasive TRKN threat. Future work will integrate improved occurrence data with climate change projections to assess likely spread pathways, building on the baseline established here.

## ACKNOWLEDGEMENTS

This work was funded by the European Union under grant agreement no. 101083727 (“NEM-EMERGE: An integrated set of novel approaches to counter the emergence and proliferation of invasive and virulent soil-borne nematodes”) and UK Research and Innovation (UKRI) under the UK government’s Horizon Europe funding guarantee grant number 10080598. In addition, BS and SS were supported by Slovenian Research and Innovation Agency (ARIS) grant P4-0072.

## AUTHOR CONTRIBUTIONS

DB, HH and SS conceived and designed the study. MD collected and curated the data and performed preliminary analyses. DB performed the final data analysis and produced the figures. MD drafted initial versions of the Methods and Results sections and produced the bivariate maps. DB wrote the final version of the manuscript. All authors contributed to manuscript review and critical discussion. All authors reviewed and approved the final version.

