## Supplementary material for "Global distributions and emergence of six major tropical root-knot nematodes"

### Supplementary Tables

**Table S1. Reported observations of *M. luci*.** Reports are listed in order of sampling date, subject to uncertainties.
Note: S = Suspected. All historical samples displaying the EST L3 esterase phenotype have been reclassified as *M. luci* (Carneiro et al., 2014)

| Location | Latitude | Sample year | Publication year | Setting | Note | References |
|---|---|---|---|---|---|---|
| Argentina | ~38˚S | ≤1985 | 2014 | NA | S | (Carneiro et al., 2014; Esbenshade and Triantaphyllou, 1985) |
| Bolivia | ~16˚S | ≤1985 | 2014 | NA | S | (Carneiro et al., 2014; Esbenshade and Triantaphyllou, 1985) |
| Ecuador | ~2˚S | ≤1985 | 2014 | NA | S | (Carneiro et al., 2014; Esbenshade and Triantaphyllou, 1985) |
| Türkiye | ~39˚N | ≤1985 | 2014 | NA | S | (Carneiro et al., 2014; Esbenshade and Triantaphyllou, 1985) |
| Caxias do Sul, RS, Brazil | 29.2˚S | ≤2000 | 2014 | Field | | (Carneiro et al., 2014, 2000) |
| Dornberk, Slovenia | 45.9˚N | 2003 | 2017 | Greenhouse | | (Gerič Stare et al., 2017; Širca et al., 2004) 25/02/2026 18:42:00 |
| Federal District of Brazil | 15.8˚S | ≤2007 | 2014 | Field | | (Carneiro et al., 2014, 2008) |
| Casablanca, Chile | 33.3˚S | ≤2007 | 2014 | Field | | (Carneiro et al., 2014, 2007) |
| Kavala, Greece | 40.9˚N | 2009 | 2017 | Field | | (Conceição et al., 2012; Gerič Stare et al., 2017) |
| Türkiye (two locations) | ~41.2˚N | 2009 | 2017 | Greenhouse | | (Aydınlı et al., 2013; Gerič Stare et al., 2017) |
| Pontecagnano, Italy | 40.6˚N | ≤2012 | 2017 | NA | | (Gerič Stare et al., 2017; Maleita et al., 2012) |
| Araucária, PR, Brazil | 25.6˚S | 2012 | 2016 | Field | | (Machado et al., 2016) |
| Coimbra, Portugal | ~40.2˚N | 2013 | 2018 | Field | | (Maleita et al., 2018) |
| Tehran, Iran | 35.7˚N | ≤2014 | 2014 | Garden | | (Carneiro et al., 2014) |
| Guatemala | ~15.8˚N | ≤2015 | 2016 | NA | | (Janssen et al., 2016) |
| Palmeira das Missões, RS, Brazil | 27.8˚S | 2015 | 2016 | Field | | (Bellé et al., 2016) |
| Frederico Westphalen, RS, Brazil | 27.6˚S | 2018 | 2019 | NA | | (Bellé et al., 2019) |
| Azores, Portugal | ~37.7˚N | 2017-2022 | 2021 | Field | | (Rusinque et al., 2021) |
| Vojvodina Province, Serbia | ~45.3˚N | 2021 | 2023 | Greenhouse | | (Bačić et al., 2023) |
| Minjar District, Ethiopia | 8.9˚N | 2021 | 2025 | Field | | (Kefelegn et al., 2025) |

**Table S2. Geospatial data sources.**

| Dataset | Variables | Resolution | Period | URL | Reference |
|---|---|---|---|---|---|
| Global maps of soil temperature | Soil temperature bioclim | 1/120° | 1979–2013 | https://zenodo.org/records/7134169 | (Lembrechts et al., 2022) |
| CHELSA V2.1 | Precipitation bioclim | 1/120° | 1981–2010 | https://www.chelsa-climate.org/datasets/chelsa_bioclim | (Brun et al., 2022) |
| GLAD | Cropland area (global overview) | 1/40° | 2019 | https://glad.umd.edu/dataset/croplands/ | (Potapov et al., 2022) |
| SoilGrids250m | Soil texture and chemistry | 1/120° | N/A | https://files.isric.org/soilgrids/former/2017-03-10/aggregated/1km/ | (Hengl et al., 2017) |
| MapSPAM V2 | Area of different crops | 1/12° | 2020 | https://doi.org/10.7910/DVN/SWPENT | (IFPRI, 2024) |
| MIRCA-OS | Crop calendars | 1/12° | 2000–2015 | https://doi.org/10.4211/hs.60a890eb841c460192c03bb590687145 | (Kebede et al., 2025) |

**Table S3. Environmental predictor VIF selection. SBIO3 was omitted due to data quality issues.**

| Code | Predictor | VIF | Included |
|---|---|---|---|
| SBIO1 | Soil annual mean temperature | 7.3 | Yes |
| SBIO2 | Soil mean diurnal range | 2.3 | Yes |
| SBIO3 | Soil isothermality | NA | No |
| SBIO4 | Soil temperature seasonality | 4.3 | Yes |
| SBIO5 | Soil maximum temperature of warmest month | >10 | No |
| SBIO6 | Soil minimum temperature of coldest month | >10 | No |
| SBIO7 | Soil temperature annual range (SBIO5-SBIO6) | >10 | No |
| SBIO8 | Soil mean temperature of wettest quarter | 2.8 | Yes |
| SBIO9 | Soil mean temperature of driest quarter | >10 | No |
| SBIO10 | Soil mean temperature of warmest quarter | >10 | No |
| SBIO11 | Soil mean temperature of coldest quarter | >10 | No |
| BIO12 | Annual precipitation | 7.0 | Yes |
| BIO13 | Precipitation of wettest month | >10 | No |
| BIO14 | Precipitation of driest month | 3.2 | Yes |
| BIO15 | Precipitation seasonality | 3.7 | Yes |
| BIO16 | Precipitation of wettest quarter | >10 | No |
| BIO17 | Precipitation of driest quarter | >10 | No |
| BIO18 | Precipitation of warmest quarter | 3.5 | Yes |
| BIO19 | Precipitation of coldest quarter | 2.2 | Yes |
| AWC | Available water capacity | 4.0 | Yes |
| BD | Bulk density of fine earth fraction | >10 | No |
| CEC | Cation exchange capacity | 3.7 | Yes |
| Clay | Clay fraction | 2.4 | Yes |
| OCC | Organic carbon content (g $kg^{-1}$) | 3.6 | Yes |
| pH | pH in water | 3.7 | Yes |
| Sand | Sand fraction | >10 | No |
| Silt | Silt fraction | 2.6 | Yes |

**Table S4. SDM algorithm performance assessed by five-fold cross validation**. Values are means across replicates for each species, the last row showing the mean of species means. Algorithms are ordered L–R by declining mean performance across species. Models were Random Forests (RF), MaxEnt, Support Vector Machines (SVM), Multiple Adaptive Regression Splines (MARS), Generalized Additive Models (GAM), Boosted Regression Trees (BRT) and Generalized Linear Models (GLM).

**a. True Skill Statistic (TSS).**

| Species | RF | MaxEnt | SVM | MARS | GAM | BRT | GLM |
|---|---|---|---|---|---|---|---|
| *M. arenaria* | 0.81 | 0.71 | 0.75 | 0.67 | 0.71 | 0.64 | 0.53 |
| *M. enterolobii* | 0.7 | 0.61 | 0.56 | 0.54 | 0.58 | 0.5 | 0.46 |
| *M. ethiopica* | 0.86 | 0.89 | 0.82 | 0.81 | 0.69 | 0.89 | 0.77 |
| *M. incognita* | 0.77 | 0.69 | 0.64 | 0.66 | 0.71 | 0.61 | 0.52 |
| *M. javanica* | 0.74 | 0.73 | 0.69 | 0.67 | 0.7 | 0.64 | 0.59 |
| *M. luci* | 0.86 | 0.87 | 0.84 | 0.86 | 0.76 | 0.82 | 0.75 |
| Mean | 0.79 | 0.75 | 0.72 | 0.7 | 0.69 | 0.68 | 0.6 |

**b. AUC**

| Method | RF | MaxEnt | MARS | BRT | SVM | GAM | GLM |
|---|---|---|---|---|---|---|---|
| *M. arenaria* | 0.94 | 0.9 | 0.87 | 0.86 | 0.9 | 0.88 | 0.78 |
| *M. enterolobii* | 0.91 | 0.84 | 0.81 | 0.81 | 0.82 | 0.83 | 0.75 |
| *M. ethiopica* | 0.94 | 0.97 | 0.91 | 0.97 | 0.87 | 0.84 | 0.91 |
| *M. incognita* | 0.93 | 0.91 | 0.89 | 0.87 | 0.88 | 0.91 | 0.82 |
| *M. javanica* | 0.92 | 0.92 | 0.89 | 0.86 | 0.87 | 0.9 | 0.84 |
| *M. luci* | 0.93 | 0.96 | 0.94 | 0.94 | 0.94 | 0.88 | 0.87 |
| Mean | 0.93 | 0.92 | 0.89 | 0.89 | 0.88 | 0.87 | 0.83 |

**Table S5. Performance of ensemble component algorithms in 30-fold bootstrap evaluation.** Values show mean ± SD across bootstrap replicates for the three algorithms retained for ensemble modelling. Values are the true skill statistic (TSS) and area under the receiver-operator curve (AUC), averaged by species.

| | TSS | | | AUC | | |
|---|---|---|---|---|---|---|
| Species | MARS | MaxEnt | SVM | MARS | MaxEnt | SVM |
| *M. arenaria* | 0.62±0.06 | 0.68±0.05 | 0.68±0.07 | 0.85±0.03 | 0.88±0.03 | 0.88±0.04 |
| *M. enterolobii* | 0.53±0.06 | 0.59±0.05 | 0.54±0.06 | 0.81±0.04 | 0.84±0.03 | 0.81±0.04 |
| *M. ethiopica* | 0.69±0.19 | 0.85±0.09 | 0.82±0.10 | 0.85±0.12 | 0.96±0.03 | 0.92±0.07 |
| *M. incognita* | 0.64±0.03 | 0.68±0.02 | 0.63±0.04 | 0.88±0.01 | 0.91±0.01 | 0.88±0.02 |
| *M. javanica* | 0.65±0.03 | 0.71±0.03 | 0.67±0.03 | 0.89±0.02 | 0.92±0.01 | 0.87±0.01 |
| *M. luci* | 0.81±0.08 | 0.82±0.08 | 0.79±0.08 | 0.93±0.05 | 0.94±0.04 | 0.92±0.05 |

**Table S6. TSS-based ensemble weights.**

| Species | MARS | MaxEnt | SVM |
|---|---|---|---|
| *M. arenaria* | 0.311 | 0.346 | 0.342 |
| *M. enterolobii* | 0.321 | 0.354 | 0.326 |
| *M. ethiopica* | 0.295 | 0.359 | 0.346 |
| *M. incognita* | 0.328 | 0.350 | 0.322 |
| *M. javanica* | 0.320 | 0.348 | 0.332 |
| *M. luci* | 0.333 | 0.340 | 0.327 |

**Table S7. Minimum training presence (MTP) and 10th-percentile training presence (P10) thresholds.**

| Species | MTP | P10 |
|---|---|---|
| *M. arenaria* | 0.020 | 0.137 |
| *M. enterolobii* | 0.022 | 0.083 |
| *M. ethiopica* | 0.044 | 0.188 |
| *M. incognita* | 0.016 | 0.085 |
| *M. javanica* | 0.016 | 0.067 |
| *M. luci* | 0.0065 | 0.126 |

**Table S8. Basal temperature ($T_b$) and lifecycle GDD sum ($S_l$) for target RKN species on different experimental host plants**. Estimates for two temperate species, the PCN *Globodera rostochiensis* and *G. pallida*, are given for comparison.

| Species | Host | $T_b$ | $S_l$ | Source |
|---|---|---|---|---|
| *G. pallida* | *Solanum tuberosum* | 4.0 | 450 | (Ebrahimi et al., 2014) |
| *G. rostochiensis* | *Solanum tuberosum* | 5.9 | 579 | (Mimee et al., 2015) |
| *G. rostochiensis* | *Solanum tuberosum* | 6.0 | 398 | (Ebrahimi et al., 2014) |
| *M. enterolobii* | *Solanum lycopersicum* | 10.0 | 507 | (Velloso et al., 2022) |
| *M. incognita* | *Phaseolus vulgaris* | 11.1 | 476 | (Giné et al., 2021) |
| *M. incognita* | *Cucumis sativus* | 11.4 | 500 | (Giné et al., 2014) |
| *M. incognita* | *Solanum lycopersicum* | 10.1 | 400 | (Ploeg and Maris, 1999) |
| *M. incognita* | *Cucumis pepo* | 12.0 | 455 | (Vela et al., 2014) |
| *M. incognita* | *Solanum lycopersicum* | 10.0 | 552 | (Velloso et al., 2022) |
| *M. incognita* | *Cucumis melo* | 14.0 | 500 | (López-Gómez et al., 2014) |
| *M. javanica* | *Cucumis sativus* | 11.4 | 500 | (Giné et al., 2014) |
| *M. javanica* | *Solanum lycopersicum* | 12.9 | 350 | (Madulu and Trudgill, 1994) |
| *M. javanica* | *Cucumis pepo* | 10.8 | 526 | (Vela et al., 2014) |
| *M. javanica* | *Cucumis melo* | 17.2 | 357 | (López-Gómez et al., 2014) |

**Table S9. Mean Galling Index (GI) and Reproductive Factor (RF) per crop**. Calculated from values reported in the literature (Supplementary Data S1). Values for all focal RKN species were combined. SPAM is the crop code used by MAPSPAM. Area is the fraction of all crop area reported by FAOSTAT for the year 2000.

| Crop | SPAM | Species | GI | RF | Area |
|---|---|---|---|---|---|
| Bananas | BANA | *Musa* spp. | 4.73 | 16.31 | 0.004 |
| Barley | BARL | *Hordeum vulgare* | 1.11 | 0.52 | 0.004 |
| Bean | BEAN | Various | 4.00 | 31.11 | 0.024 |
| Cassava | CASS | *Manihot esculenta* | 3.15 | 0.90 | 0.021 |
| Citrus | CITR | *Citrus* spp. | 0.00 | 0.00 | 0.003 |
| Coconut | CNUT | *Cocos nucifera* | 0.00 | 0.04 | 0.008 |
| Coffee (arabica) | COFF | *Coffea arabica* | 4.28 | 32.67 | Note |
| Coffee (robusta) | RCOF | *Coffea robusta* | 2.00 | 1.62 | 0.008 |
| Cotton | COTT | *Gossypium* spp. | 2.46 | 17.85 | 0.022 |
| Cowpeas | COWP | *Vigna unguiculata* | 0.63 | 0.76 | 0.011 |
| Groundnuts (Peanuts) | GROU | *Arachis hypogaea* | 3.13 | 1.13 | 0.021 |
| Maize | MAIZ | *Zea mays* | 3.25 | 7.02 | 0.138 |
| Millet | MILL | *Panicum miliaceum*, etc. | 0.15 | 1.03 | 0.022 |
| Pigeon Pea | PIGE | *Cajanus cajan* | 2.42 | 17.36 | 0.004 |
| Potato | POTA | *Solanum tuberosum* | 3.60 | 56.73 | 0.012 |
| Rapeseed | RAPE | *Brassica rapa* | 2.12 | 12.06 | 0.024 |
| Rice | RICE | *Oryza sativa* | 2.75 | 2.06 | 0.114 |
| Sorghum | SORG | *Sorghum bicolor* | 0.82 | 0.95 | 0.028 |
| Soybeans | SOYB | *Glycine max* | 2.99 | 13.395 | 0.088 |
| Sugar beet | SUGB | *Beta vulgaris* | 5.00 | 15.50 | 0.003 |
| Sugarcane | SUGC | *Saccharum officinarum* | 1.88 | 2.80 | 0.018 |
| Sunflower | SUNF | *Helianthus annuus* | 2.45 | 23.09 | 0.019 |
| Sweet Potato | SWPO | *Ipomoea batatas* | 4.04 | 16.84 | 0.005 |
| Tomatoes | TOMA | *Solanum lycopersicum* | 3.20 | 46.40 | 0.004 |
| Wheat | WHEA | *Triticum aestivum* | 2.04 | 3.30 | 0.15 |
| Yams | YAMS | *Discorea* spp. | 2.25 | 0.90 | 0.007 |

Note: FAOSTAT does not distinguish between Arabica and Robusta coffee, the value for all coffee is given in the Robusta row.

### Supplementary Figures

**Figure S1. Observed distributions of RKN species**. Points indicate the locations of RKN species compiled from the scientific literature and other sources. Shading indicates cropland.

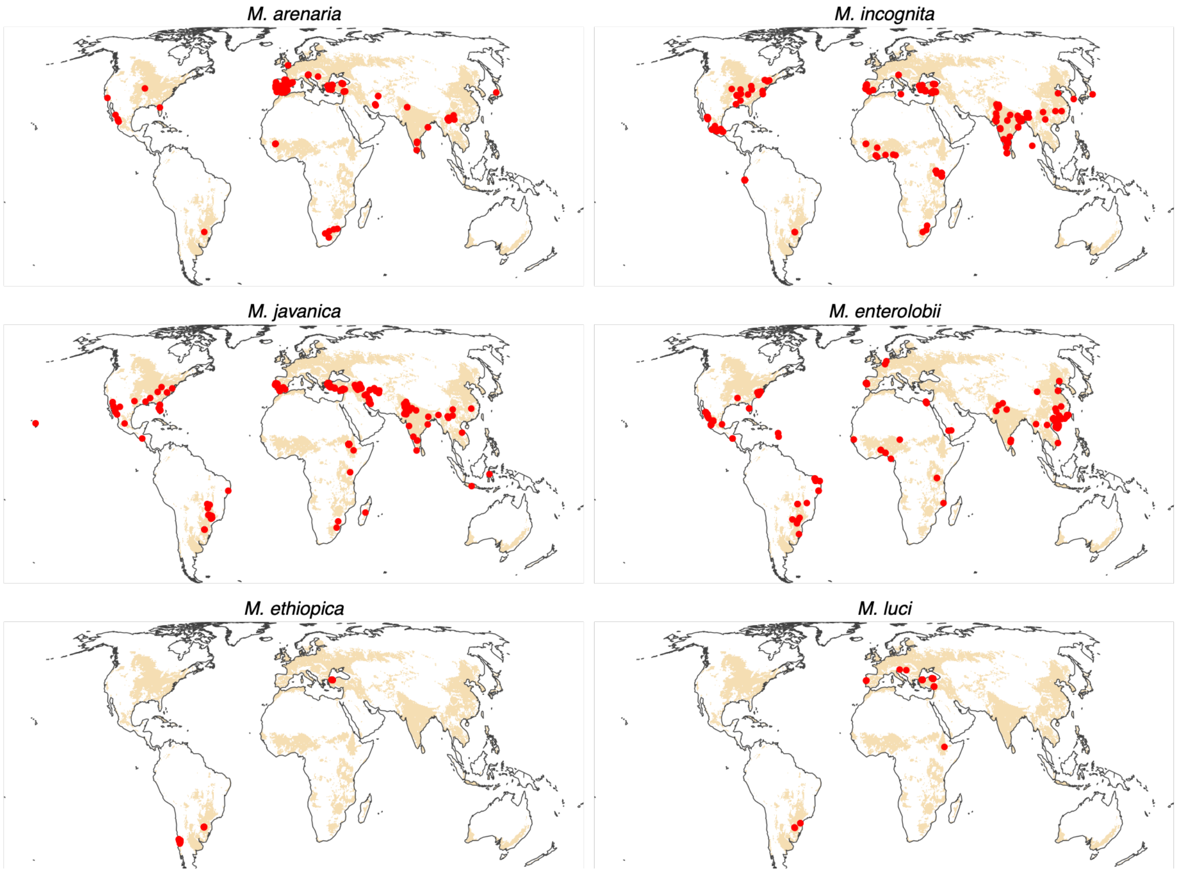

**Figure S2. Predictor variable importance**. Boxplots show the distribution (median, interquartile range, and range) of variable importance values across 30 bootstrap replicates. Importance was calculated as the absolute reduction in model AUC following random permutation of each predictor.

(Figure on next page)

(Caption on previous page)

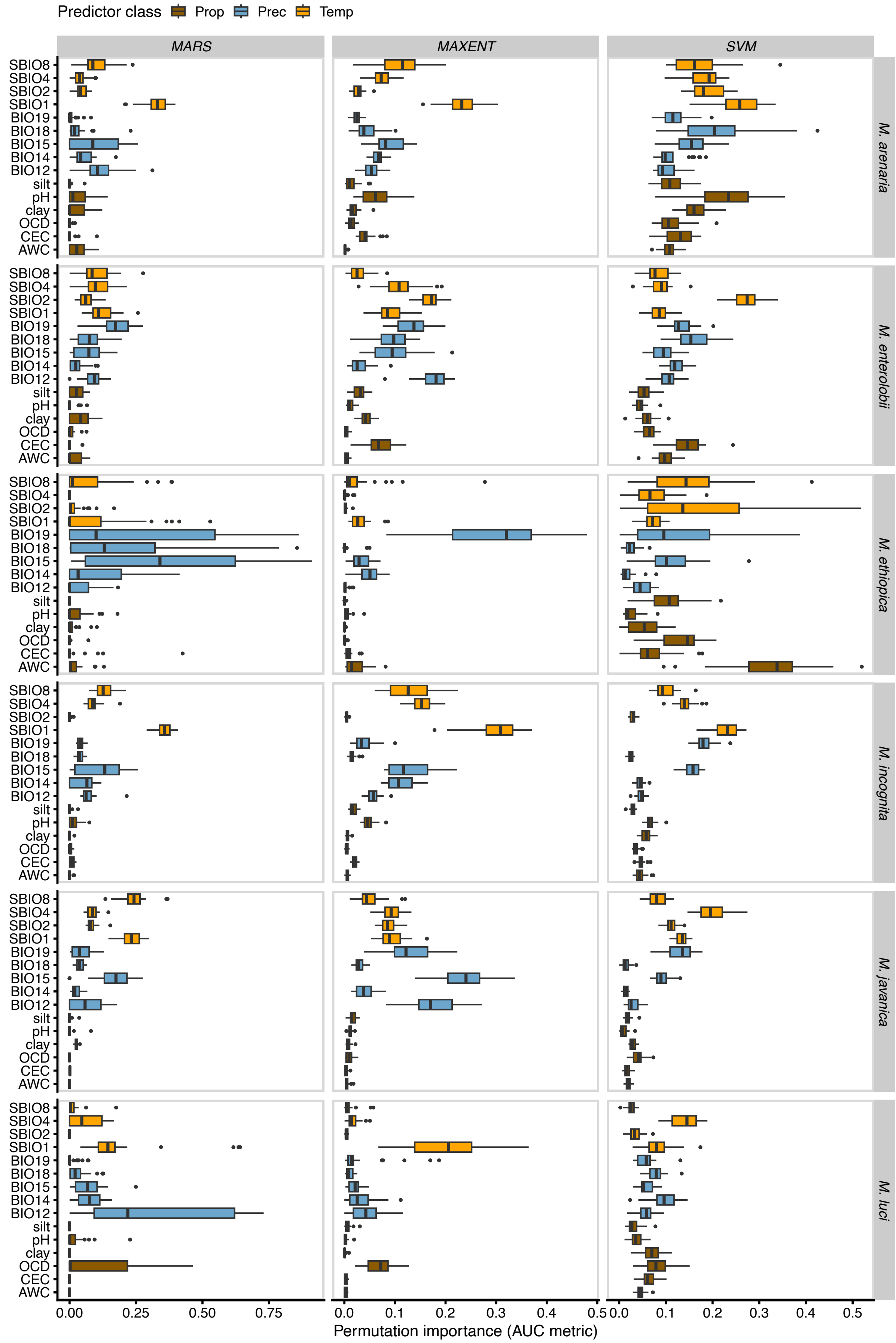

**Figure S3. Suitability response curves**. Ensemble response curves (thick black lines) are bootstrap mean TSS-weighted means of individual model response curves (thin coloured lines). Predictors are ordered alphabetically. Response curves for each species are shown on separate pages.

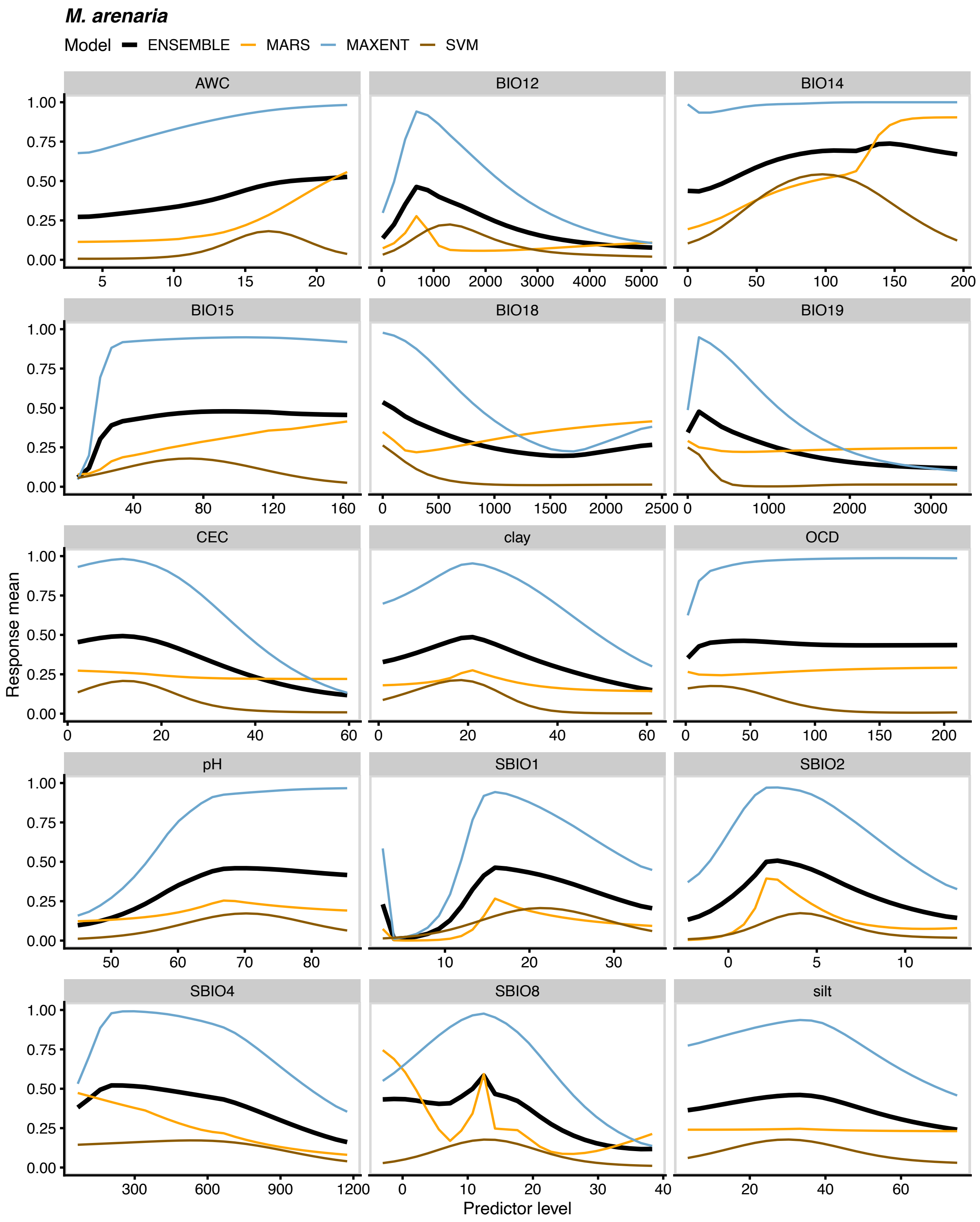

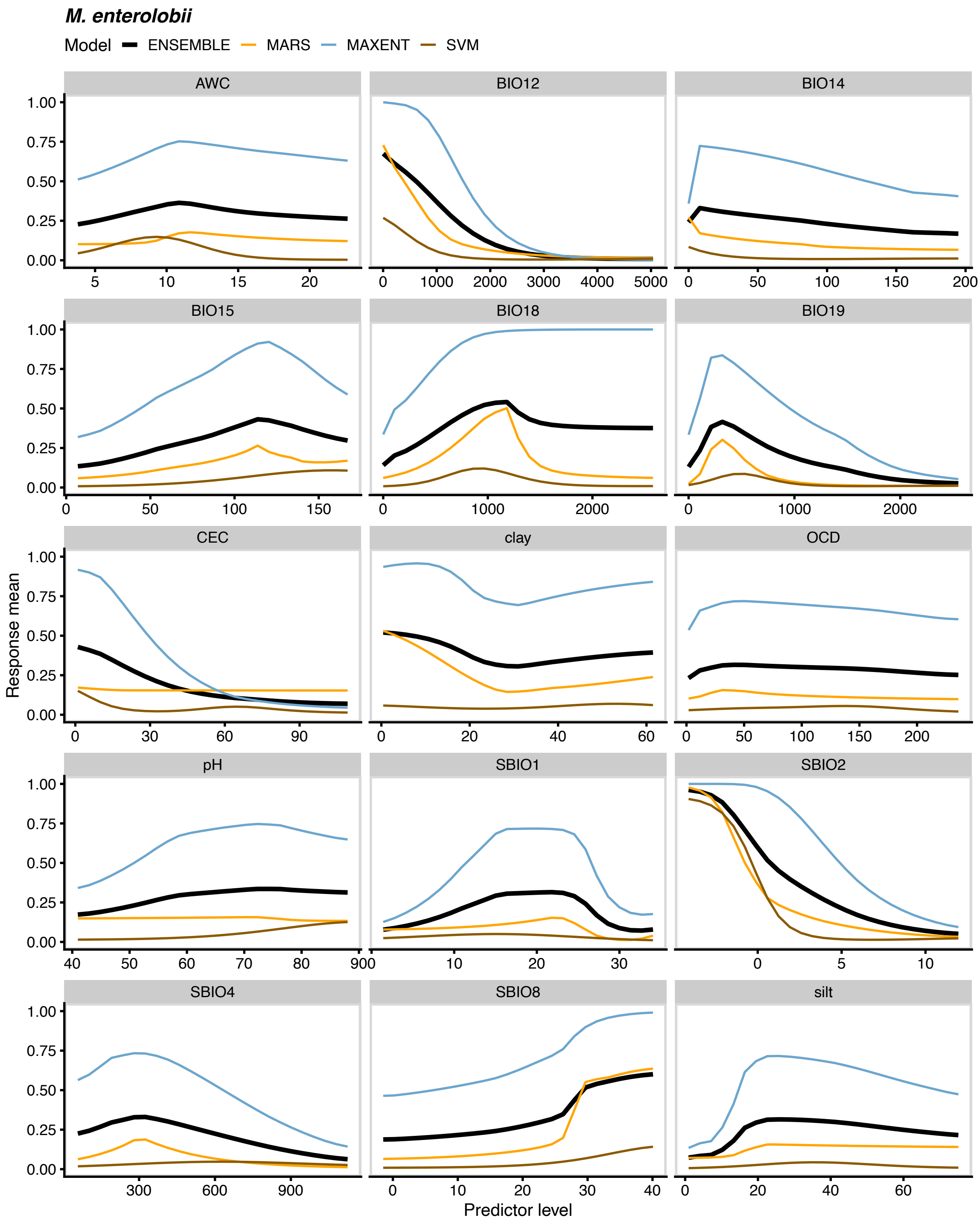
M. enterolobii
Model ENSEMBLE MARS MAXENT SVM
AWC
BIO12
BIO14
BIO15
BIO18
BIO19
CEC
clay
OCD
pH
SBIO1
SBIO2
SBIO4
SBIO8
silt
Response mean
Predictor level
0.00 0.25 0.50 0.75 1.00

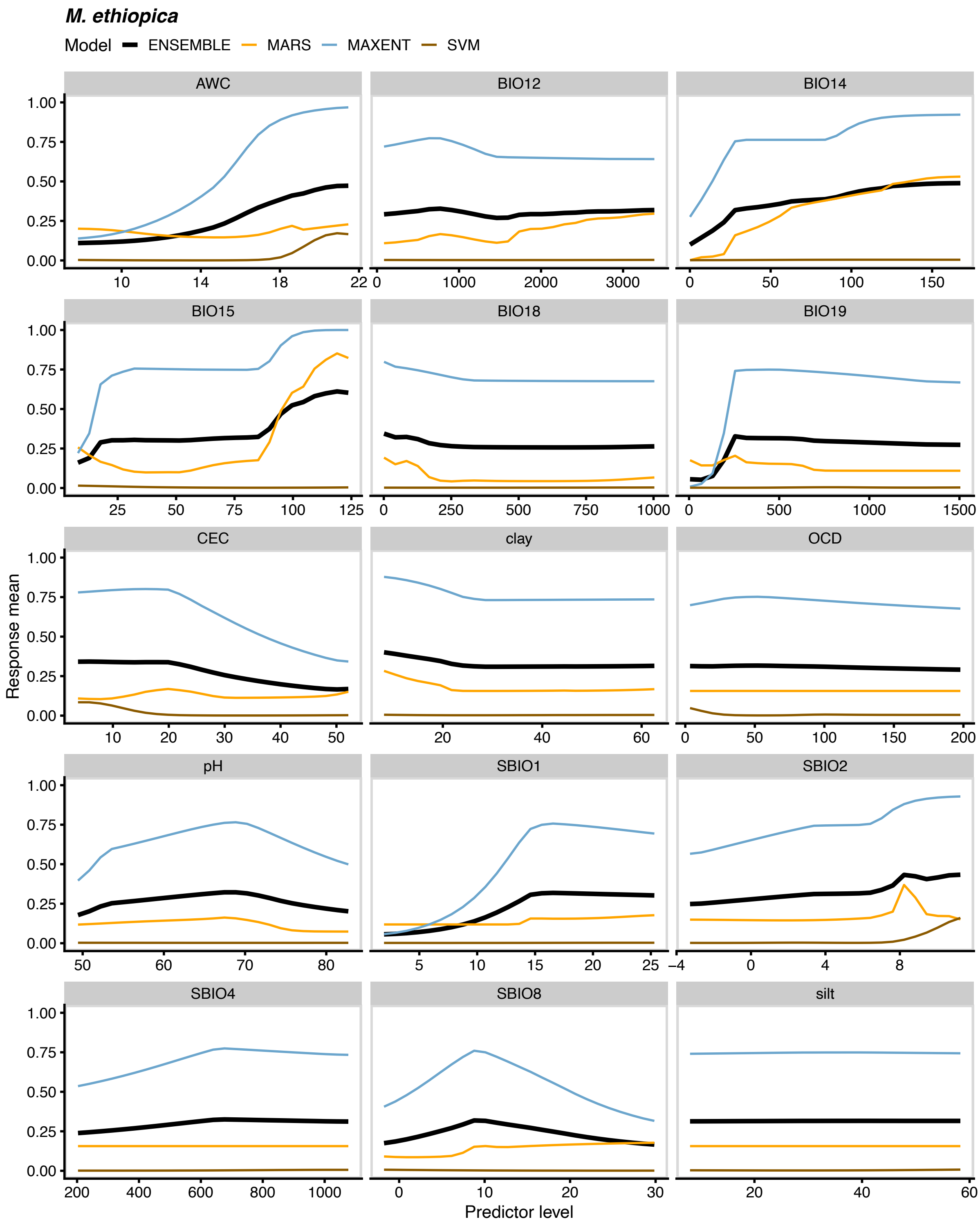
M. ethiopica
Model ENSEMBLE MARS MAXENT SVM
AWC
BIO12
BIO14
BIO15
BIO18
BIO19
CEC
clay
OCD
pH
SBIO1
SBIO2
SBIO4
SBIO8
silt
Response mean
Predictor level
1.00
0.75
0.50
0.25
0.00

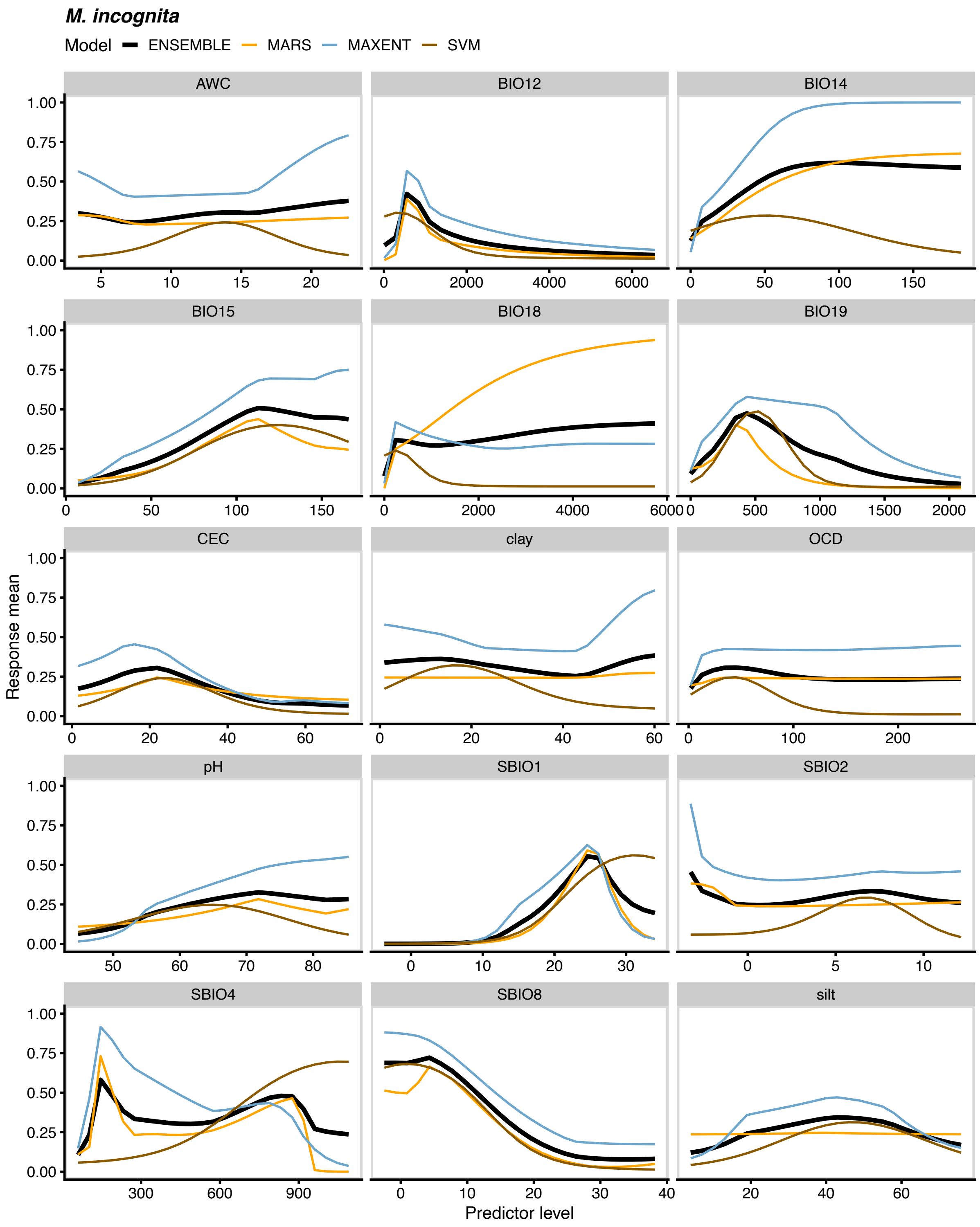
M. incognita
Model ENSEMBLE MARS MAXENT SVM
AWC
BIO12
BIO14
BIO15
BIO18
BIO19
CEC
clay
OCD
pH
SBIO1
SBIO2
SBIO4
SBIO8
silt
Response mean
Predictor level

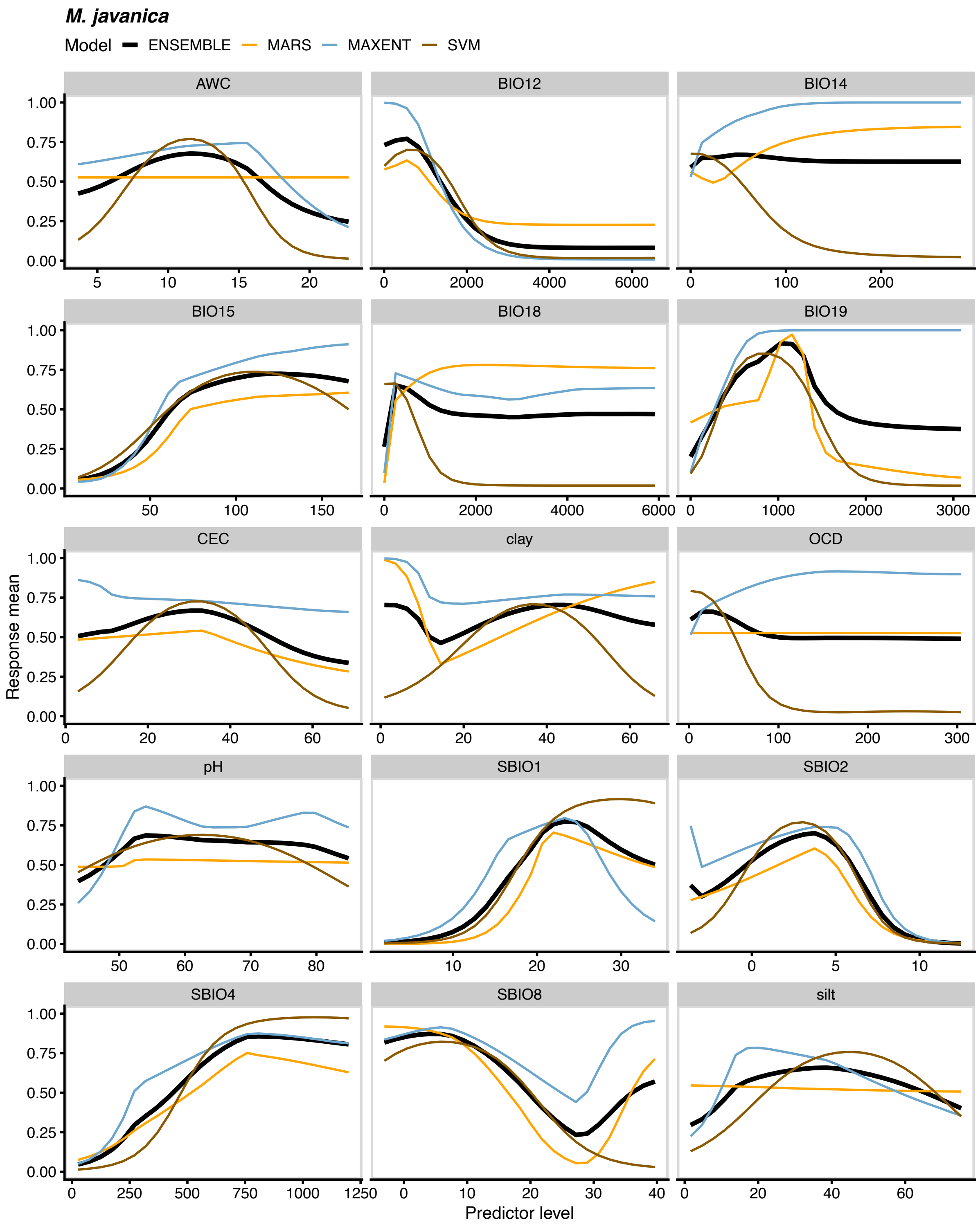
M. javanica
Model ENSEMBLE MARS MAXENT SVM
AWC
BIO12
BIO14
BIO15
BIO18
BIO19
CEC
clay
OCD
pH
SBIO1
SBIO2
SBIO4
SBIO8
silt
Response mean
Predictor level

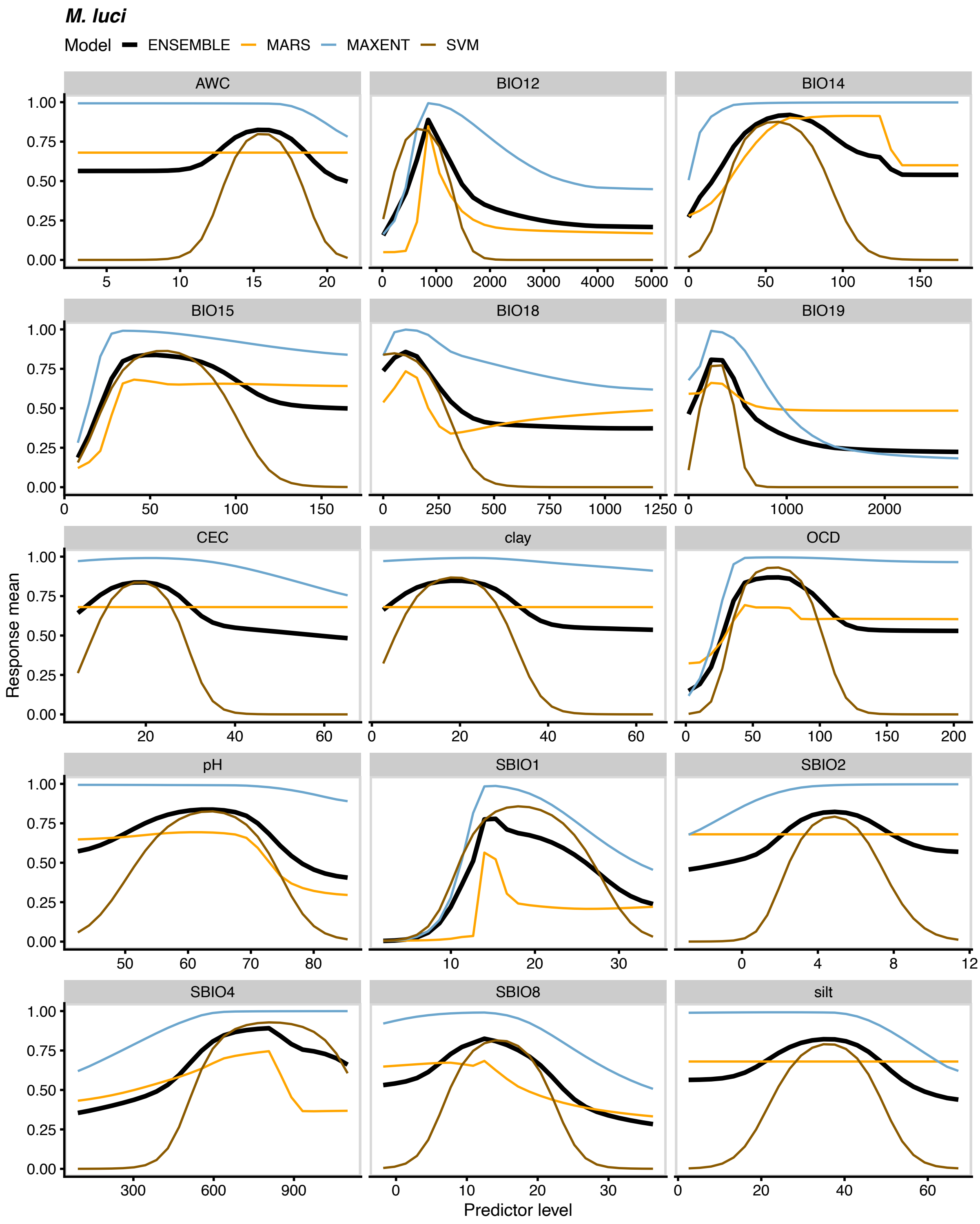

M. luci
Model ENSEMBLE MARS MAXENT SVM
AWC
BIO12
BIO14
BIO15
BIO18
BIO19
CEC
clay
OCD
pH
SBIO1
SBIO2
SBIO4
SBIO8
silt
Response mean
Predictor level

**Figure S4. Ensemble habitat suitability.** Values are bootstrap mean TSS-weighted averages of MARS, MaxEnt and SVM model predictions.

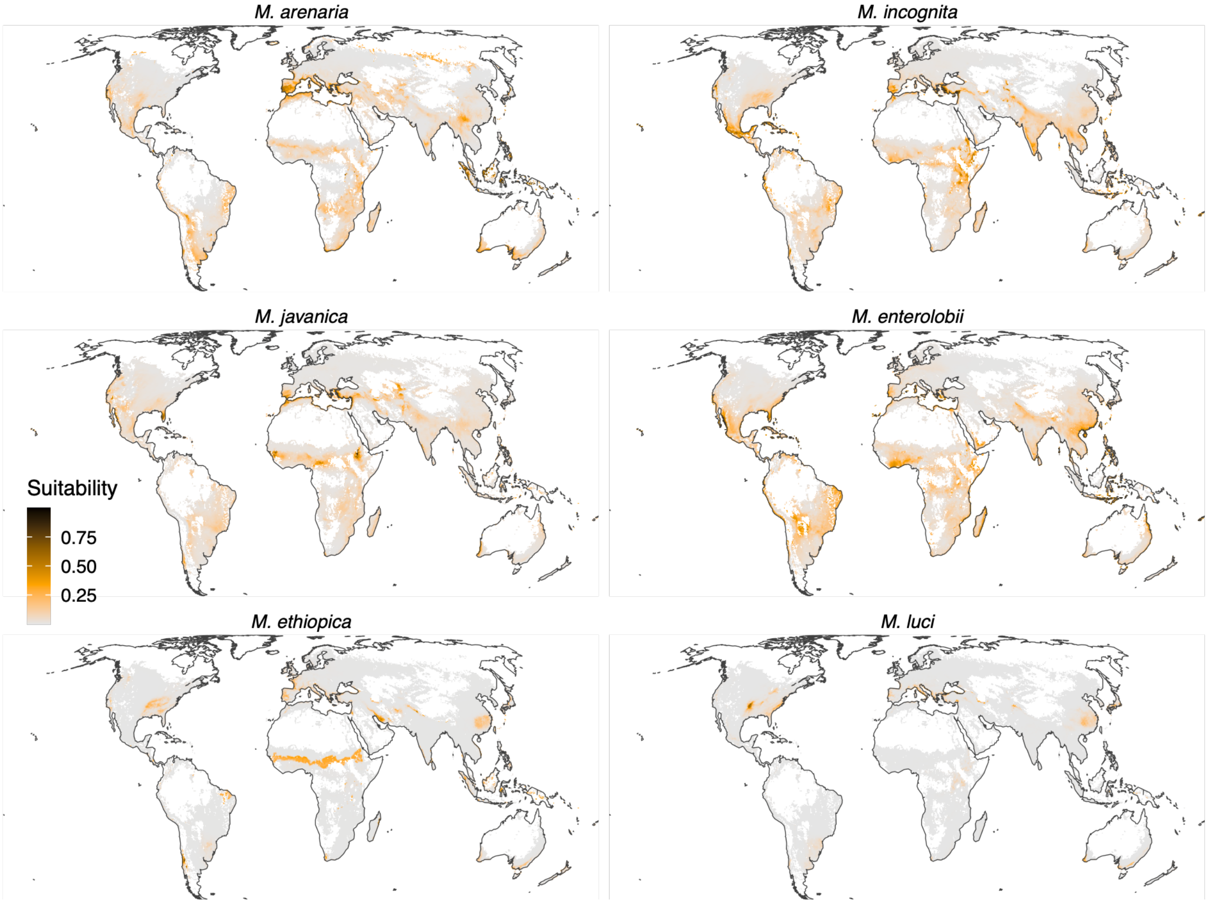

**Figure S5. Combined suitability for six RKN species, Europe.** Colours indicate the number of species predicted to occur under the MTP threshold (grey indicating zero).

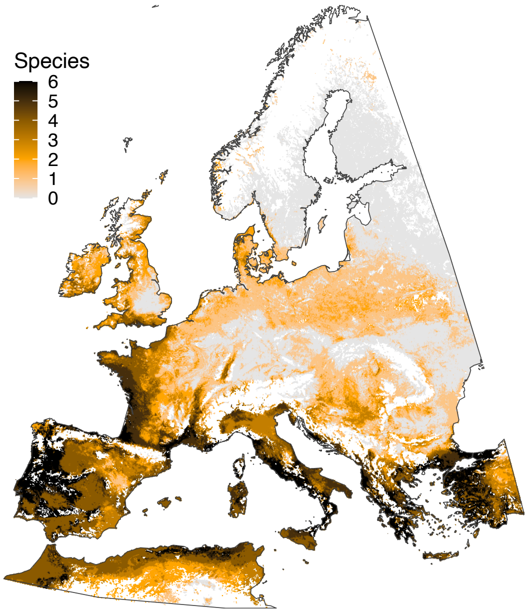

**Figure S6. Life cycle GDD sums ($S_l$) and base temperatures ($T_b$) for life-cycle completion**. Three RKN (orange symbols) compared with two potato cyst nematodes (*Globodera* spp.). Data extracted from the literature (see Supplementary Table S2).

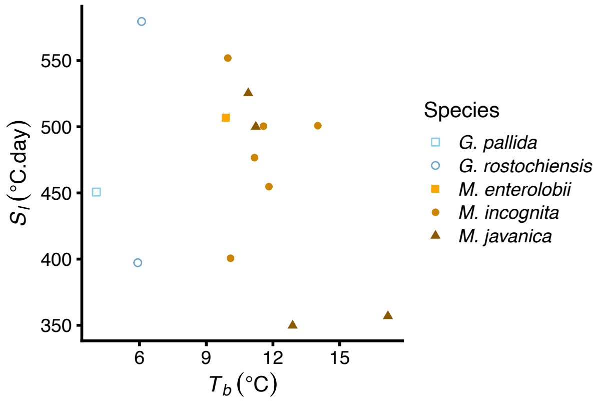

**Figure S7. Distributions of predicted number of generations at 0-5 cm soil depth**. Estimates are from various studies of life-cycle degree day requirements for three RKN species grown on different hosts. Hosts are melon (*Cucumis melo*), pumpkin (*Cucurbita pepo*), cucumber (*Cucumis sativus*), common bean (*Phaseolus vulgaris*) and tomato (*Solanum lycopersicum*). Boxplots show median (thick line), interquartile range (box) and outliers (whiskers) of a random sample of $10^5$ global cropland grid cells (0.025˚ × 0.025˚). Colours indicate studies (Giné et al., 2021, 2014; López-Gómez et al., 2014; Madulu and Trudgill, 1994; Ploeg and Maris, 1999; Vela et al., 2014; Velloso et al., 2022).

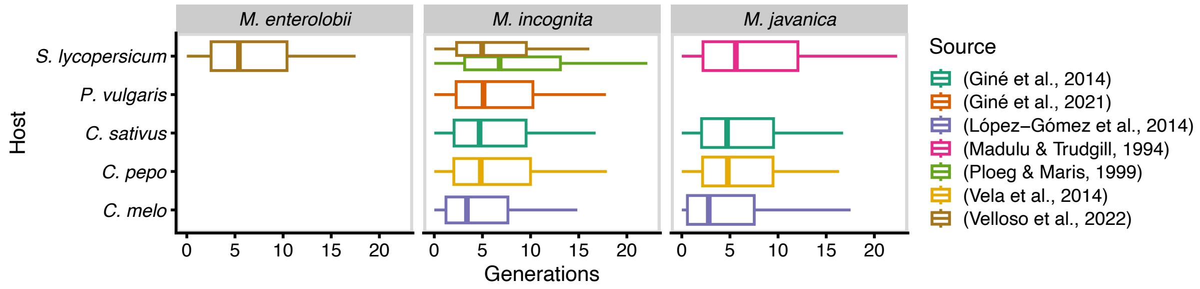


**Figure S8. Difference between mean predicted number of generations per year estimated from air and soil temperatures**. a) Maps of differences. b) Histograms of differences. Predictions were restricted to global cropland.

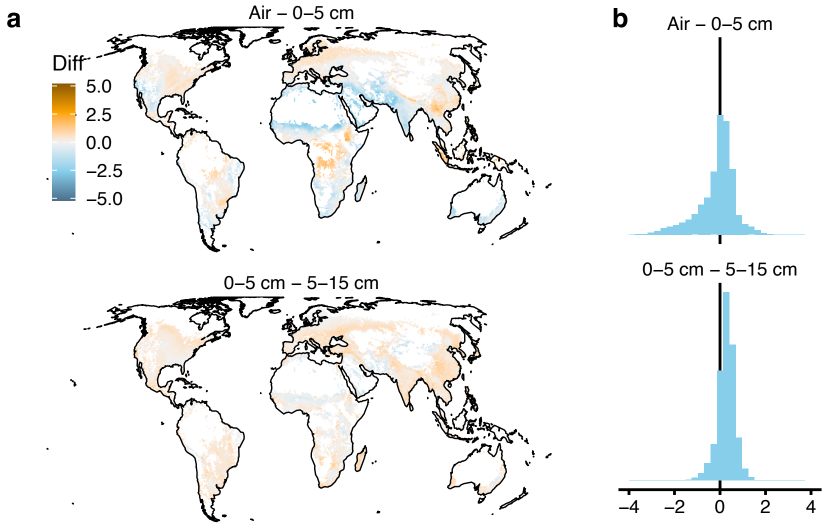

**Figure S9. TRKN generations per year**. a) Mean life cycles per year estimated from mean monthly air temperatures and soil temperatures at 0–5 cm and 5–15 cm, calculated from eleven $S_l$ and $T_b$ estimates for three TRKN species. b) Standard deviations of mean across 11 estimates.

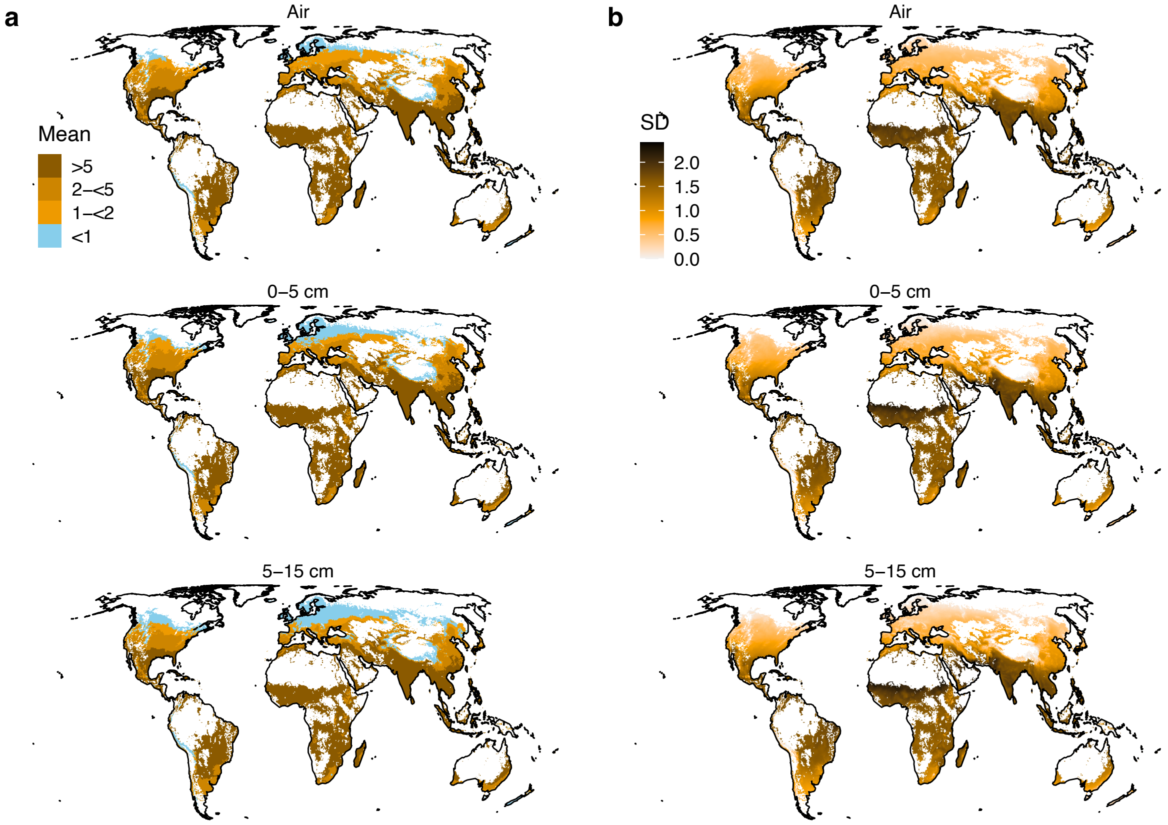

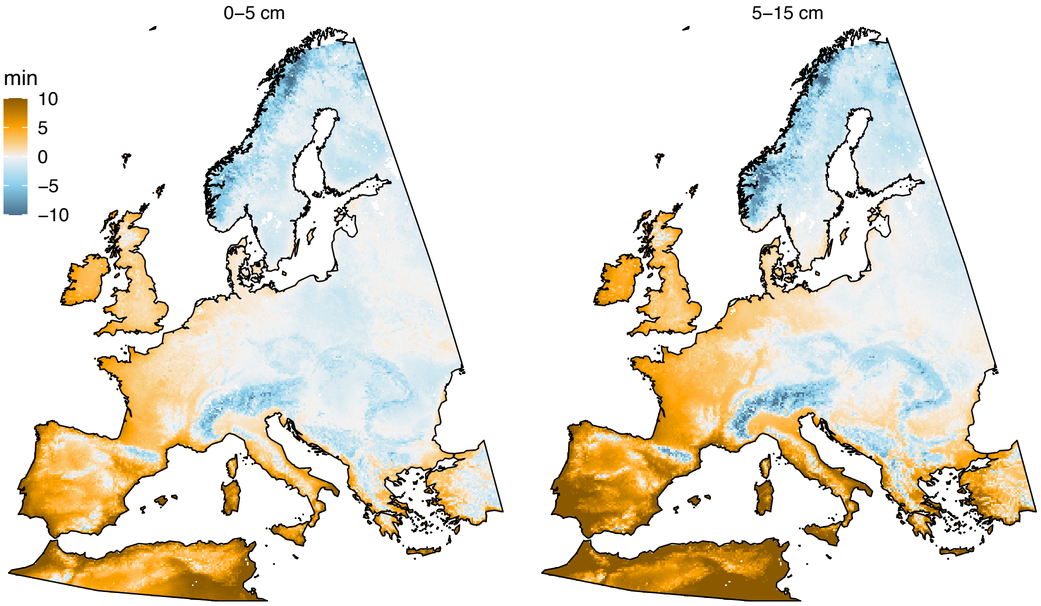


**Figure S10. Minimum temperature of coldest month (BIO6) in Europe**. The colour scale is constrained to the range –10 °C to 10 °C, with values outside this range shown in the nearest category.

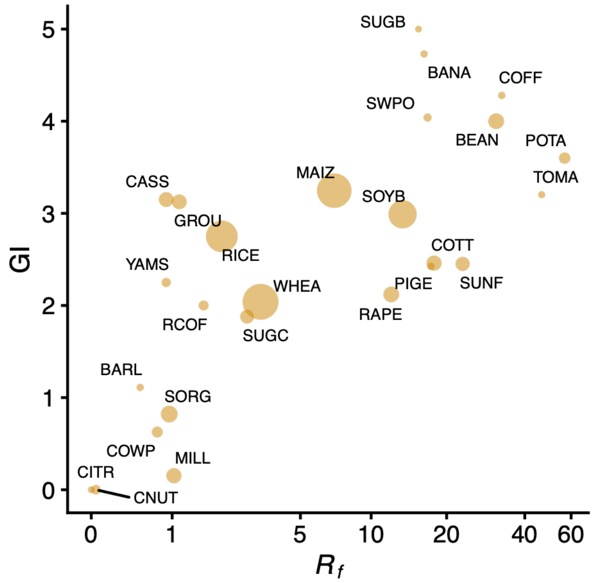


**Figure S11. Galling Index (GI) vs reproductive factor (RF) for major crops**. Values represent means per crop across values reported in the literature (Supplementary Data S2). Labels are crop codes from the MAPSPAM dataset (Supplementary Table S8). Symbol sizes indicate relative area of production reported in FAOSTAT for the year 2000 (Supplementary Table S8), except Arabica coffee (COFF) which is arbitrary as this crop is not evaluated independently in FAOSTAT.

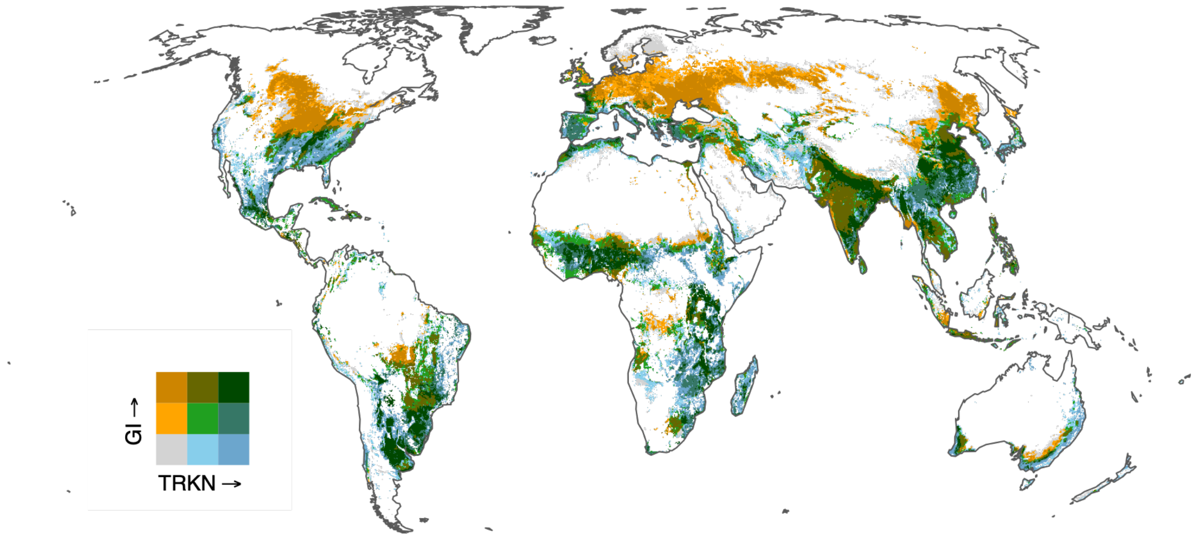


**Figure S12. Combined TRKN habitat and crop suitability**. Habitat suitability is based on the sum of MTP-threshold SDM predictions, and crop suitability on the crop area-weighted galling index (GI). Colour scales were derived from quantiles of habitat suitability and GI. Blue indicates high habitat but low crop suitability, orange indicates low habitat but high crop suitability, and green indicates regions where both are suitable for TRKN proliferation. Grey denotes areas unsuitable for both.

### Interactive Figures

Interactive versions of Figure 6 and Supplementary Figure S12 are available at:
https://mohammeddakhil1.github.io/NEM-EMERGE-Interactive-Bivariate-Map_RF/
https://mohammeddakhil1.github.io/NEM-EMERGE-Interactive-Bivariate-Map_GI/
